# Biology System Description Language (BiSDL): a modeling language for the design of multicellular synthetic biological systems

**DOI:** 10.1101/2024.01.13.575499

**Authors:** Leonardo Giannantoni, Roberta Bardini, Alessandro Savino, Stefano Di Carlo

## Abstract

**Background:** The Biology System Description Language (BiSDL) is an accessible, easy-to-use computational language for multicellular synthetic biology. It allows synthetic biologists to represent spatiality and multi-level cellular dynamics inherent to multicellular designs, filling a gap in the state of the art. Developed for designing and simulating spatial, multicellular synthetic biological systems, BiSDL integrates high-level conceptual design with detailed low-level modeling, fostering collaboration in the Design-Build-Test-Learn cycle. BiSDL descriptions directly compile into Nets-Within-Nets (NWNs) models, offering a unique approach to spatial and hierarchical modeling in biological systems.

**Results:** BiSDL’s effectiveness is showcased through three case studies on complex multicellular systems: a bacterial consortium, a synthetic morphogen system and a conjugative plasmid transfer process. These studies highlight the BiSDL proficiency in representing spatial interactions and multi-level cellular dynamics. The language facilitates the compilation of conceptual designs into detailed, simulatable models, leveraging the NWNs formalism. This enables intuitive modeling of complex biological systems, making advanced computational tools more accessible to a broader range of researchers.

**Conclusions:** BiSDL represents a significant step forward in computational languages for synthetic biology, providing a sophisticated yet user-friendly tool for designing and simulating complex biological systems with an emphasis on spatiality and cellular dynamics. Its introduction has the potential to transform research and development in synthetic biology, allowing for deeper insights and novel applications in understanding and manipulating multicellular systems.

## I. Introduction

Computational methods play a crucial role in synthetic biology, providing powerful tools that significantly improve the design, analysis, and construction of synthetic biological systems, with a particular emphasis on multicellular synthetic systems [1], [2]. These systems implement intricate functions by distributing genetic constructs among different cells [3]. This approach exploits intra and intercellular interactions within the cell population, distributing the metabolic burden to amplify system responsiveness. However, the complexity of these synthetic designs leads to intricate interactions with the host organism, thereby diminishing predictability and controllability [4], [5]. In this context, synthetic morphogenesis applications pose unique challenges as they strive to govern cellular self-organization, which heavily relies on spatial relationships and interactions among cells in space [6].

Computational methods have a key role in the analysis [7], [8], modeling [9], [10], design [1] and optimization [11], [12], [13] of complex biological processes. In particular, for analyzing and predicting the dynamics of multicellular synthetic systems, computational tools must offer instruments for modeling and simulation, accounting for multiple spatial and temporal scales [14]. Computational modeling languages serve as powerful tools in this domain. They must expressively represent the target systems while integrating knowledge from diverse sources [15], thus enhancing our understanding of the Design–Build–Test–Learn (DBTL) cycle [16]. Furthermore, to facilitate interdisciplinary collaboration, these languages must support collaborative development, reproducibility, and knowledge sharing [17], [18].

In computational biology, Domain-Specific Languages (DSLs) serve specific applications (see section II). For instance, the Systems Biology Markup Language (SBML) specializes in biochemical networks [19], NeuroML focuses on the structure and function of neural systems [20], and the Simulation Experiment Description Markup Language (SED-ML) handles procedures for running computational simulations [21]. While these languages excel within their applications, their limited scope and interoperability [22] hamper integration into the multi-level models needed for multicellular synthetic biology. The Infobiotics Language (IBL) addresses interoperability by consolidating modeling, verification, and compilation into a single file, streamlining *in silico* synthetic biology processes and ensuring compatibility with the Synthetic Biology Open Language (SBOL) and SBML frameworks [22]. However, IBL lacks support for describing multicellular synthetic designs and expressing spatial aspects crucial for synthetic morphogenesis applications [6].

Models of multicellular, spatial biological systems can utilize low-level modeling formalisms [23], [24], [25] or multi-level hybrid models that combine different formalisms across multiple scales [14]. Unfortunately, these powerful tools are primarily accessible to expert users, limiting their availability to experimental synthetic biologists.

This paper introduces the Biology System Description Language (BiSDL), a computational language for spatial, multicellular synthetic designs that can be directly compiled into simulatable, low-level models to explore system behavior. BiSDL aims to balance simplicity and intuitive usage for broad accessibility, while its expressive power enables the description of biological complexity in multicellular synthetic systems. Building on preliminary work [26], BiSDL supports flexible abstraction, allowing non-experts to reuse high-level descriptions and experts to manipulate or create low-level models. Additionally, BiSDL supports modularity and composition, facilitating the creation and usage of libraries for knowledge exchange, integration, and reuse in the multicellular synthetic biology DBTL cycle. In this work, the low-level models generated from BiSDL descriptions are based on the Nets-Within-Nets (NWN) formalism [27], chosen for its capability in multi-level and spatial modeling of complex biological processes [25], [28]. Nevertheless, the language is general enough to be integrated with other low-level formalisms.

BiSDL closely mirrors the natural language used within the biological domain. The compiler manages the gap between this high-level biological semantics and the low-level NWN formalism syntax, reducing the need for advanced modeling skills and knowledge of the low-level formalism. While in its current implementation, BiSDL requires basic programming and modeling skills, the language could be integrated with a dedicated Graphical User Interface (GUI), paving the way for extensive broadening of the user base. BiSDL aims to simplify data exchange in bioinformatics, offering a high-level approach that abstracts away the complexity found in standards like SBOL [29], [30] and SBML [31], and is versatile enough to be translated into other formalisms like Petri Nets (PN). Unlike tools such as pySBOL [32] and libSBML [33] that still rely on complex XML-like syntax.

The paper is organized as follows: section II summarizes existing computational languages for synthetic biology, section III details the design, syntax, and semantics of BiSDL and its compilation into low-level models using the NWN formalism. Then, section IV showcases BiSDL capabilities through three case studies on multicellular synthetic systems. Finally, section V summarizes the contributions, highlights open challenges, and outlines future developments.

## II. Related work

The scientific landscape of model description languages for systems and synthetic biology is rich and complex.

Figure 1 organizes them by expressivity of biological semantics and generality in modeling different biological levels or domains. Each language links to the biological levels it targets (either molecular pathways, cells, multicellular systems, or a combination thereof) and the level of flexibility the language has in generalizing to different biological domains or mechanisms (see Legend on the right).

**Fig. 1.**
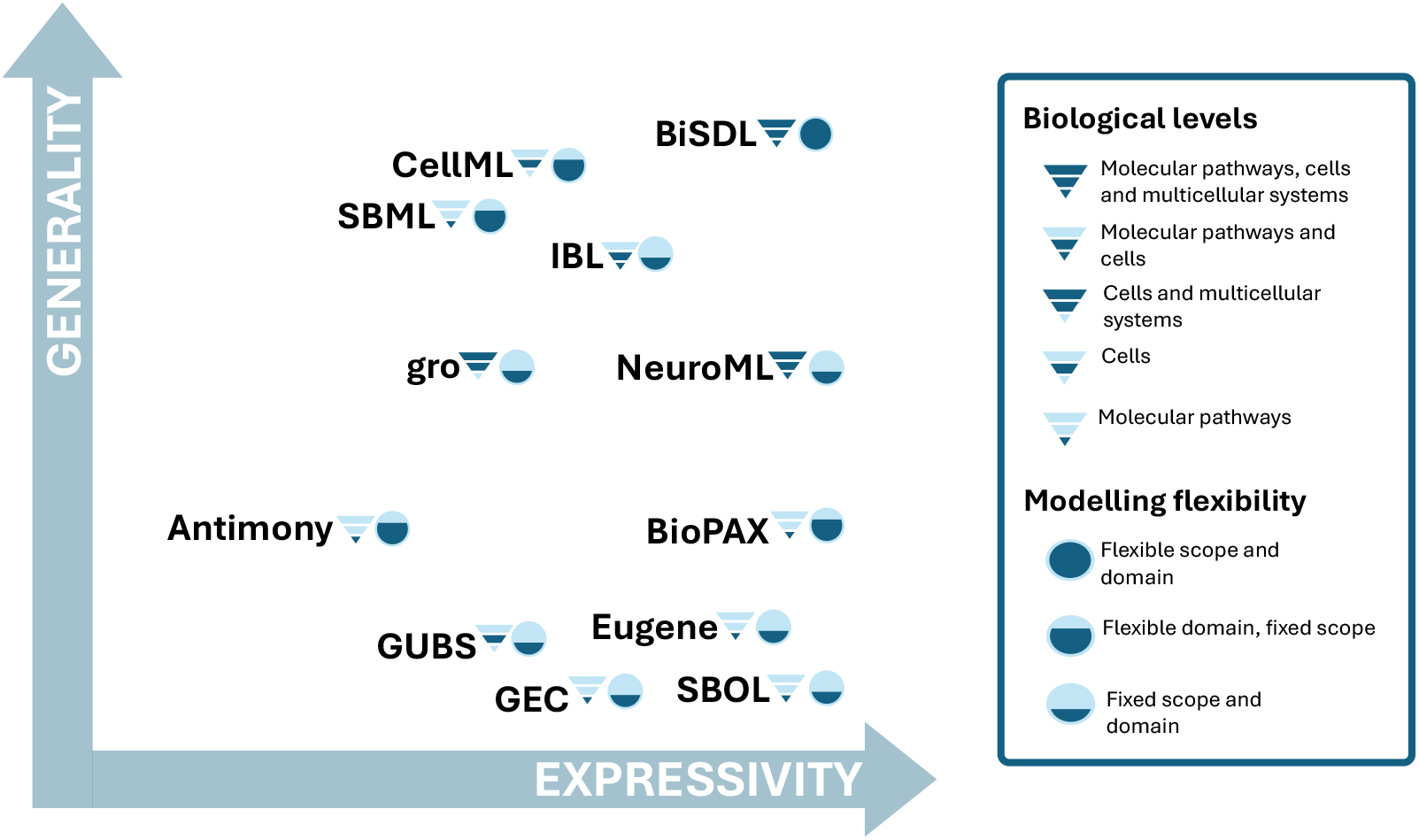
Comparison of Model Description Languages in Systems Biology over expressivity (broadness and depth of described models) and generality (broadness of modeling target, scope, and domain), providing further details on the biological levels covered and the modeling flexibility supported.

The COmputational Modelling in BIology NEtwork (COMBINE) initiative coordinates the development of inter-operable and non-overlapping standard languages covering different aspects of biological systems [34], [35], [36]. COMBINE DSLs provide intermediary layers between the user and low-level modeling formalisms. They rely on `XML` for model description and compile into Ordinary Differential Equations (ODE) models, making this mathematical modeling formalism accessible by non-expert users. Some of the COMBINE DSLs specialize in intracellular pathways, such as BioPAX [37], and processes, such as Systems Biology Graphical Notation (SBGN) [38], SBML [31] and CellML [39]. SED-ML [40], [21] exclusively aims at managing simulations of system behavior. NeuroML [20] tackles different biological aspects simultaneously, including spatiality and support simulation management, yet specializes in only neuronal systems. Among COMBINE standards, SBOL [29], [30] targets *in silico* synthetic genetic designs, yet is limited to genetic circuits alone, and does not cover any other biological aspect. Existing standards such as SBOL [29], [30], CellML [39], SED-ML [40], [21], and SBML [31] offer valuable frameworks for data exchange, albeit with limitations such as verbosity and complexity. While tools like pySBOL [32] and libSBML [33] provide programmatic access to these standards, they still require users to navigate XML-like syntax. BiSDL allows users to focus solely on high-level concepts, abstracting from the implementation details. The proposed compiler, translating BiSDL into PN, showcases the versatility of the language, demonstrating the potential for translation into other languages and formalisms.

Besides COMBINE standards, several computational languages for systems and synthetic biology exist [41]. Antimony is a text-based definition language that directly converts to the SBML standard employed in Tellurium, a modeling environment for systems and synthetic biology [42]. The Cell Programming Language (gro) [43] is a language for simulating colony growth and cell-cell communication in synthetic microbial consortia. It handles the spatiality and mobility of bacterial cells, internal genetic regulations, and mutual communications. Eugene [44] specifies synthetic biological parts, devices, and systems, mainly focusing on genetic constructs and their expression. Genetic Engineering of Cells (GEC) [45] centers over logical interactions between proteins and genes. GEC programs can be compiled into sequences of standard biological parts for laboratory applications. Genomic Unified Behavior Specification (GUBS) [46] focuses on the cell’s behavior as the central entity with a rule-based, declarative, and discrete modeling formalism. gro and GUBS can model the interaction between cells. gro also supports the representation of the spatial organization in a system. However, it is limited to bacterial cells only. Also, both languages require programming skills. Thus, neither is easily accessible to non-expert users. Eugene and GEC focus on genetic circuits only or simple molecular interaction networks for representing and exchanging reusable genetic designs through functional modules, such as Standard Parts [47]. Even when combined in more complex structures, such modules only partly comprise the complexity and hierarchy of interdependent regulations and the role of spatiality in biological multicellular designs. IBL [22] is a DSL for synthetic biology that manages several computational aspects into a single specification, overcoming interoperability issues and ensuring seamless compatibility with SBOL and SBML frameworks. Yet, it currently does not support multicellular synthetic systems nor spatial aspects.

To overcome the limitations of existing solutions in biological scope, expressivity of multicellular and spatial aspects, and accessibility, BiSDL provides high-level descriptions of intra- and inter-cellular mechanisms over spatial grids, and their direct compilation into low-level simulatable models for the exploration of system behavior.

## III. Methods

To help synthetic biologists create models of multicellular synthetic systems, BiSDL is designed for users with varying computational skills, making biological knowledge readable and writable. BiSDL stands at an abstraction level parallel to biological concepts used in experimental science, bridging the gap between these concepts and the more intricate low-level models. BiSDL descriptions combine user-friendly biological semantics with the capacity to capture system complexity. Its development is centered on domain-specific terminology and the ability to compile into NWN simulation models, discussed in subsection III-D. The language syntax covers process hierarchies, spatial relations, and cellular interactions at the intercellular and intracellular levels. While the BiSDL supports the description of the system, the BiSDL models can be compiled into complex NWN models for the simulation and analysis of system behavior.

### A. Biological perspectives and levels of abstraction

The BiSDL syntax supports describing spatial and multi-level biological concepts through multiple domains and abstraction levels, as illustrated in the Y Chart reported in Figure 2.

**Fig. 2.**
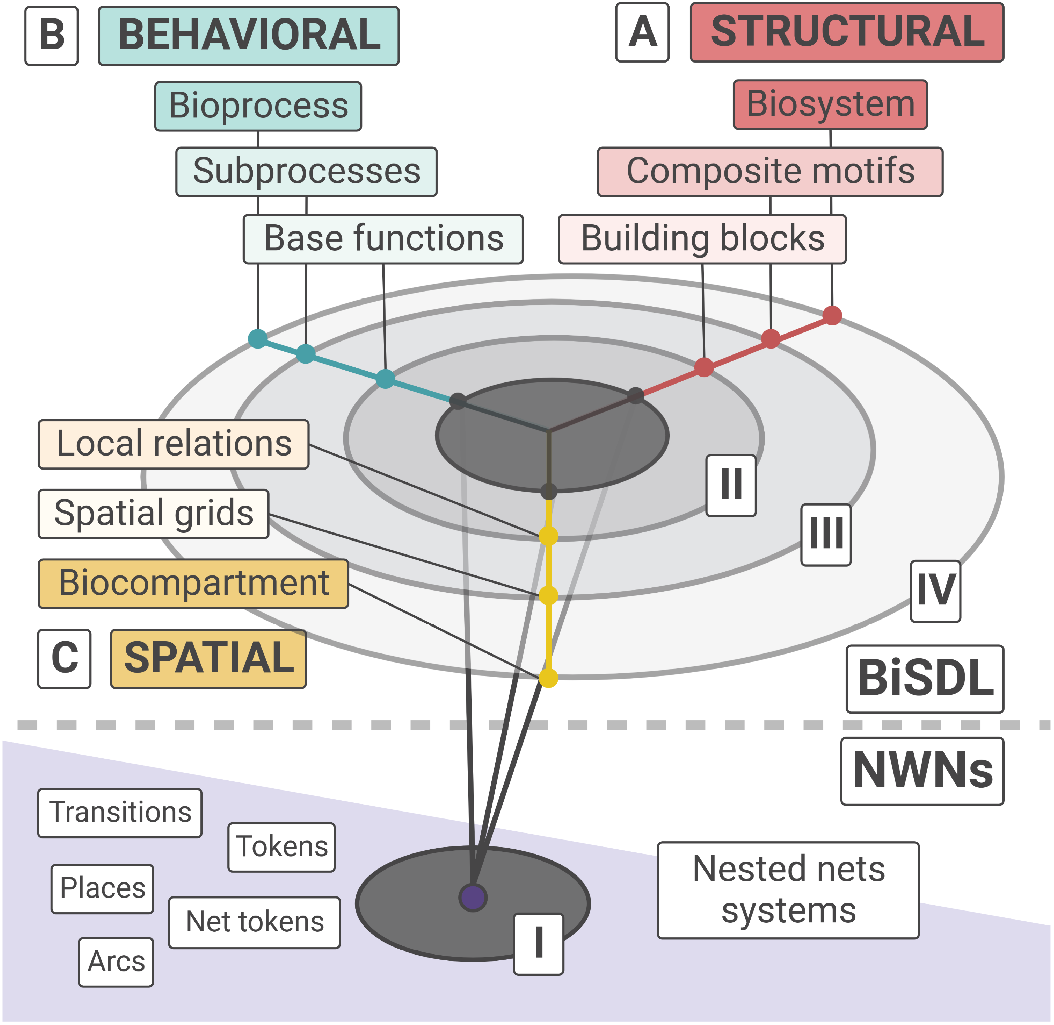
A scheme of BiSDL domains and abstraction levels inspired to the VHDL Y-Chart [48]. The upper panel (BiSDL) shows the (A) Structural, (B) Behavioral, and (C) Spatial domains and the corresponding high abstraction levels (II, III, and IV) in BiSDL descriptions. The lower panel (NWN) illustrates the three domains at the low abstraction level relative to the Nets-Within-Nets formalism (I).

Inspired by the Very High-Speed Integrated Circuit Hardware Description Language (VHDL) [48], the BiSDL Y Chart adapts the three description domains defined for VHDL (i.e., Behavioral, Structural, and Physical) to the biological semantics.

The *Structural* domain, illustrated in Figure 2 (A - STRUCTURAL, top right), delves into the architecture of biological structures such as transcriptional machinery, protein complexes, or synthetic genetic constructs within a host. On the other hand, the *Behavioral* domain (Figure 2, B - BEHAVIORAL, top left) focuses on describing the dynamic functioning, interactions, and transformations of biological elements, encompassing processes like gene transcription, diffusion, and protein degradation. Lastly, the *Spatial* domain (Figure 2 C - SPATIAL, center left) outlines the spatial substrate influencing interactions among the elements composing the system (e.g., the spatial organization of a group of cells).

The BiSDL describes each domain at four different abstraction levels, spanning from general biological concepts (high abstraction) to the NWN modeling formalism elements (low abstraction).

*Level IV* (Figure 2, circle IV) describes a *Biosystem* comprising multiple composite motifs within the structural domain. This corresponds, for instance, to a *Bioprocess* made up of multiple bioprocesses in the behavioral domain and a *Biocompartment* defined by multiple spatial grids in the spatial domain. *Level III* (Figure 2, circle III) elucidates *Composite motifs*, i.e., combinations of building blocks representing complex biological structures in the structural domain. These correspond to *Subprocesses* that emerge from interlaced base functions in the behavioral domain, necessitating *Spatial grids* in the spatial domain to model the underlying spatial relations. *Level II* (Figure 2, circle II) defines *Building blocks*, capturing fundamental biological concepts in the structural domain. These correspond to *Base functions* in the behavioral domain and simple *Local relations* in the spatial domain. *Level I* (Figure 2, circle I) describes NWN formalism elements combined in low-level models of the system.

When describing composite systems (e.g., a biological tissue), BiSDL covers all domains: structural, spatial, and behavioral (as shown in Figure 2.A-C). These descriptions can include cells, extracellular structures, spatial arrangements, and the processes involved, requiring elements from each of the BiSDL domains. However, simpler descriptions may focus on a single domain, such as the transcription process of a gene concentrating on the behavioral domain. Simple `MODULE` definitions can be combined to form more complex descriptions. The syntax and semantics of a set of composable BiSDL descriptions are detailed in Additional file 1 - Section 1 as an example of a BiSDL library.

### B. Syntax and semantics

All BiSDL descriptions respect the template shown in algorithm 1 based on a hierarchy of `MODULE`, `SCOPE` and `PROCESS` constructs. They start with naming the `MODULE` (line 1). A `MODULE` encapsulates the complete description of a biological system, encompassing structural, behavioral, and spatial aspects. This includes detailing groups of cells, their spatial arrangement on a two-dimensional grid, intracellular processes, and the spatial diffusion mechanisms facilitating intercellular communications. Modules are self-contained and serve as the fundamental units for reusing and composing existing descriptions.

Each `MODULE` consists of a set of `SCOPE` declarations with defined identifiers and spatial coordinates (lines 3-12 and 13) that describe the relevant biological compartments within the modeled system and a set of `DIFFUSION` mechanisms (lines 14-15) that model the diffusion of signals among them. The `SCOPE` declarations may incorporate additional communication methods, such as PARACRINE_SIGNAL (line 10) and `JUXTACRINE_SIGNAL`(lines 11-12), describing intercellular communication, either through diffusible signals (paracrine) or direct contact (juxtacrine). Integer timescales can represent any ratio between the operations of different models in the provided discrete-time simulator. The `TIMESCALE` of a module (line 2) sets the base pace of the system dynamics compared to the unitary step of the discrete-time simulator. For instance, if one model has a `TIMESCALE` of N, it means that it evolves by 1 step every N simulator’s steps (whose `TIMESCALE` is made equal to 1). The model is slower than the base time step by a factor of N.

Each `SCOPE` contains a set of biological `PROCESS` instantiations with explicit identifiers (lines 4-8 and 9). They comprise *base functions* like transcription, translation, and degradation.

The `TIMESCALE` of a process (line 6) is a discrete multiplier of the `MODULE` timescale, determining the relative speed at which the process occurs compared to the base module pace. The same applies to processes with different timescales: they proceed at a relative speed, the ratio of their respective timescales. For instance, if `TIMESCALE` is 2 for `PROCESS` *p1*, and 5 for `PROCESS` *p2*, they will proceed at a relative speed of 5/2 (i.e., *p1* evolves 2.5 times faster than *p2*). Different `PROCESS` instances can connect over the same elements: for example, one process might produce a molecule that regulates a base function in another process. BiSDL emphasizes ease of description. Each `SCOPE` can reuse a `PROCESS` from another `SCOPE` simply by declaring a `PROCESS` with the same `<process_id>`.

#### Algorithm 1: BiSDL general template. Each MODULE organizes around a set of SCOPE definitions. Each SCOPE contains a set of PROCESS instances describing the behavior of entities in the MODULE and a set of SIGNAL declarations describing communication mechanisms among entities. DIFFUSION mechanisms support communication among SCOPE constructs.

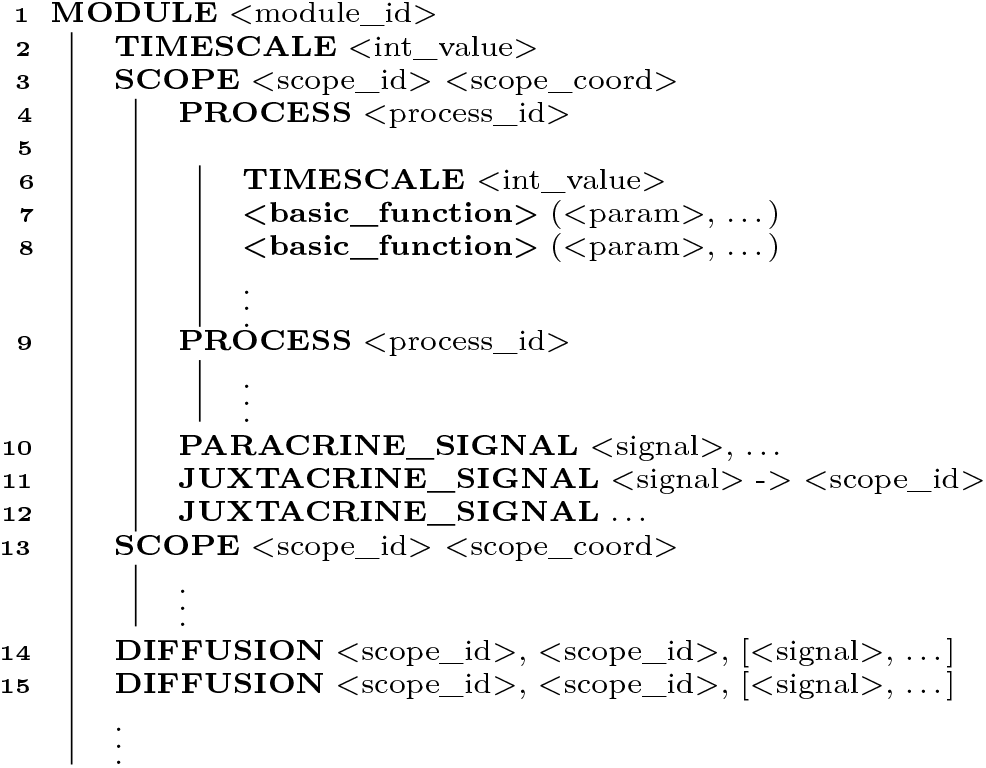

As a simple example, algorithm 2 provides the BiSDL description of the chemical reaction by which two *H*_2_ molecules react with one *O*_2_ molecule to form two *H*_2_*O* molecules.

#### Algorithm 2: BiSDL description of the chemical reaction by which two *H*_2_ molecules react with one *O*_2_ molecule to form two *H*_2_*O* molecules.

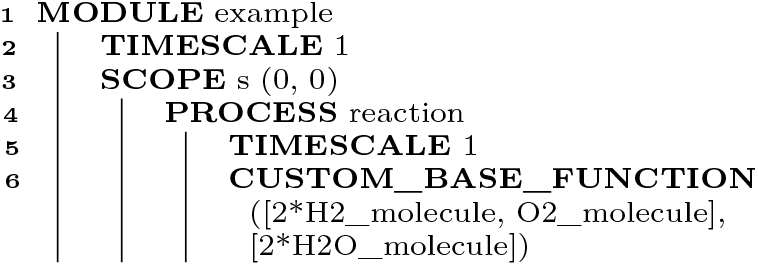

The `MODULE`, whose base `TIMESCALE` is `1`, consists of a biological compartment at coordinates `(0, 0)` within the spatial grid (`SCOPE s`, line 3). The `SCOPE` contains a single `PROCESS`, whose base `TIMESCALE` multiplier is `1`, named `reaction`. Here, the entities describing molecular hydrogen (`H2_molecule`) and oxygen (`O2_molecule`) transform into water (`2*H2O_molecule`, line 6). Multipliers for `H2_molecule` and `H2O_molecule` specify the proportion the molecules combine, implying a multiplier with unitary value when not indicated.

### C. BiSDL constructs

This work proposes a library of BiSDL constructs (see Additional file 1 - Section 1) to exemplify the language semantic capabilities, showing its expressiveness and closeness to the biological semantics. To foster standardization in model description languages, all proposed BiSDL constructs follow the Systems Biology Ontology (SBO) [49] and fall into the following four subcategories (of the seven provided by the standard):

- Physical entity representation:
  – Material entities identify the functional entities (i.e., `SCOPE`, `CELL`, and the base types `GENE`, `MRNA`, `PROTEIN`, `COMPLEX`, `MOLECULE`);
  – Functional entities identify the function they perform (i.e., PARACRINE_SIGNAL, `JUXTACRINE_SIGNAL`, and `DIFFUSION`).
- Participant role:
  – identifies the role played by an entity in a modeled process (i.e., `INDUCERS`, `INHIBITORS`, `ACTIVATORS`);
- Occurring entity representation:
  – identifies processual relationships involving physical entities (i.e., `TRANSCRIPTION`, `TRANSLATION`, `DEGRADATION`,`PROTEIN_COMPLEX_FORMATION`, `ENZYMATIC_REACTION`, `CUSTOM_PROCESS`);
- System description parameter:
  – provides quantitative descriptions of biological processes (i.e., `TIMESCALE`s and the multipliers of physical entities).

### D. From BiSDL descriptions to NWNs models

BiSDL supports the system dynamics analysis utilizing simulations. This is obtained by compiling BiSDL descriptions into low-level models based on the NWN formalism.

#### 1) Compilation of NWNs models

NWN extend the PN formalism to support hierarchy, encapsulation, and selective communication [27], [24], [9], which makes them suitable to model complex biological processes [25], [28], coherently with the design goals of BiSDL (subsection III-A). PN are bipartite graphs where nodes can be either *places* or *transitions*. Places represent states the modeled resources can assume. Transitions model the creation, consumption, or transformation of resources in, from, or across places. Tokens model discrete units of resources in different states. Each transition has rules regulating its enabling and activation, depending on the availability of tokens in its input places. Directed arcs link places and transitions to form the desired network architectures. When a transition fires, it consumes the required tokens from the input places and creates tokens in its output places.

The NWN formalism is a high-level PN formalism supporting all features of other high-level PN: tokens of different types and timed and stochastic time delays associated with transitions. NWN introduce an additional type of token named *Net Token*. A Net Token is a token that embeds another instance of a PN. With this type of token, NWN support hierarchical organization, and each layer relies on the same formalism (see Figure 3). This characteristic introduces the Object-Oriented Programming (OOP) paradigm within the PN formalism. Therefore, NWN models express encapsulation and selective communication, allowing the representation of biological compartmentalization and semi-permeability of biological membranes easily. Nets at different levels in the hierarchy evolve independently and optionally communicate through synchronous channels that interlock transitions from different nets, synchronizing their activation upon satisfaction of enabling conditions.

**Fig. 3.**
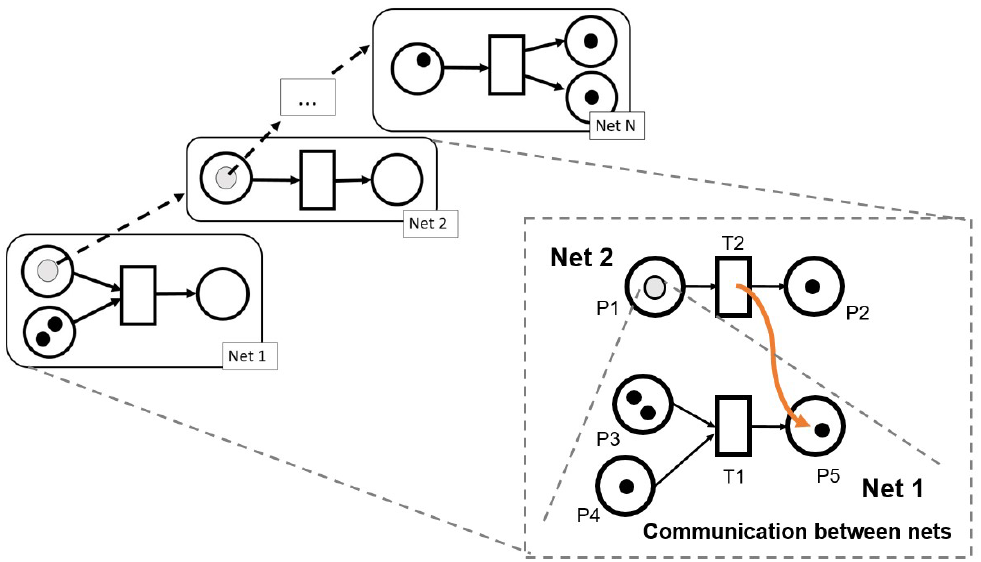
A representation of the NWN formalism. Tokens in these PN can be instances of PN, thus implementing a hierarchy of encapsulated levels. Channels can interlock two transitions from different nets, allowing the exchange of tokens and information.

NWN have heightened expressivity compared to other modeling formalisms. While Boolean models offer binary node states, high-level Petri Nets can convey intricate information regarding system resources and processes. Similarly, while ODE represent uniform compartments with continuous values, NWN accommodate discrete and continuous quantities. Unlike many existing approaches that primarily focus on intracellular mechanisms, multilevel NWN allow for the modeling of both intracellular and supra-cellular information, enabling a broader scope of representation and effective analysis of complex biological systems [23]. However, the increased expressivity of NWNs comes at the cost of greater model complexity and computational demands for simulation algorithms, a common trade-off in computational modeling [50]. In conclusion, the decision to employ NWN for demonstrating BiSDL stems from the desire to highlight its full expressive capacity in generating complex models. BiSDL may support compilation into various low-level formalisms, generating models on different points across this trade-off.

To support NWN, BiSDL compilation generates models implemented with `nwn-snakes`, a customized version of the `SNAKES` library [51]. `SNAKES` is an efficient Python library for the design and simulation of PN [51]. The `nwn-snakes` library presented in this work extends `SNAKES` to handle multi-scale models, ensuring consistency across the hierarchical levels in the model. `nwn-snakes` provides constructs to express the hierarchy of temporal and spatial scales. Every BiSDL `MODULE` in the compiled model is represented by a `Module` class with an individual timescale, which in turn inherits from the `PetriNet` class implemented in `nwn-snakes`. A prototype BiSDL compiler (`bisdl2snakes.py`) generates Python `Module` classes implementing `nwn-snakes` models from BiSDL descriptions. Detailed instructions on compiler use are available in the BiSDL `GitHub` public repository (see Availability of data and materials).

The spatial hierarchy underlying BiSDL descriptions (see subsection III-B) is translated into a low-level model based on a system of nested spatial grids represented by PN. In this model, the places model sub-portions of space, allowing the representation of multiple spatial scales. Each place in a spatial grid can host, as a net token, another spatial grid, ensuring cross-level semantic consistency across different spatial scales. This work provides consistent semantics for two-level hierarchies, which support the intended modeling of multicellular systems where both intra- and intercellular mechanisms are described. `nwn-snakes` handles the marking evolution of the two levels synchronously: if marking evolves on one level, the other level mirrors the exact change. Additional file 1 - Section 2 reports the way `nwn-snakes` supports NWN modeling describing the mapping between BiSDL building blocks and NWN.

BiSDL supports high interpretability of generated NWN models in two ways. Firstly, a compilation of BiSDL constructs labels the resulting low-level constructs with the high-level specific parameters. For instance, the construct `PROTEIN_COMPLEX_FORMATION(3*LuxR_protein, 3*AHL_molecule, 3*LuxR_AHL_complex)` generates NWN constructs containing the product name: `LuxR_AHL_complex`. Secondly, in BiSDL, any construct can be wrapped into a process, and the process is assigned a custom name. This feature supports the direct reuse of processes in general and the reuse of constructs wrapped up in processes by leveraging the process name. algorithm 4 exemplifies this mechanism: the `PROCESS` defined in lines 18-21 encapsulates `TRANSCRIPTION`, `TRANSLATION`, and `DEGRADATION` constructs, and is named `CD19_production`. The same process is reused (by name) in line 37. Indeed, the `SCOPE` defined in lines 36-40 (5 lines of code) reuses processes defined only once for the first `SCOPE`, which spans over 35 lines of code (1-35). In compilation, to avoid ambiguity, each time the same process is used again in the BiSDL description, a new set of low-level elements is generated and named by appending a progressive number. In the same example from Algorithm 4, the first instance of `PROCESS CD19_production` (lines 18-21) is assigned the name `CD19_production_process_0` in the NWN model. The second instance (line 37) is internally assigned the name `CD19_production_process_1`, and thereafter.

#### 2) Simulation of NWNs models

The exploration of systems dynamics relies on the simulation of `nwn-snakes` models compiled from BiSDL descriptions with the `nwn-petrisim` simulator. The simulator is designed to be simple and easy to use, thus requiring minimal coding as reported in Listing 1.

**Listing 1.**
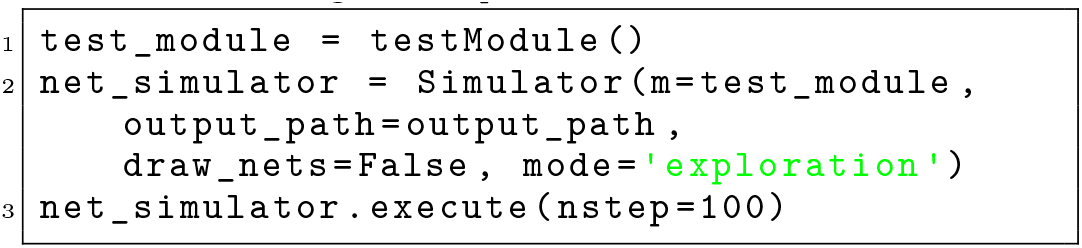
`nwn-petrisim` instantiation

The simulator is instantiated at line 2. The argument `m=test_module` represents the instance of the top-level net to be simulated. The arguments `draw_nets=False` and `mode=‘exploration’` control the generation of visual output, preventing the creation of images of the net architectures and allowing evolution plots to adapt to the generated output value ranges.

The simulation, executed in line 3, is discrete, with `nstep=100` determining the number of simulation steps. The simulator analyzes the stochastic evolution of the system. Additionally, it allows simulating the system’s response to external stimuli applied to the model as outlined in Listing 2. The simulation comprises a loop running for the specified number of steps (`n_steps`). Within this loop, at every `n` steps, the simulator adjusts the marking of the place that models the stimulus within the network. This adjustment involves adding a random number of black tokens, ranging from 0 to `r`. Subsequently, this modified marking is applied to the simulated model before proceeding to the next simulation step. The values of `n` and `r` control the intensity of the administered stimulus.

**Listing 2.**
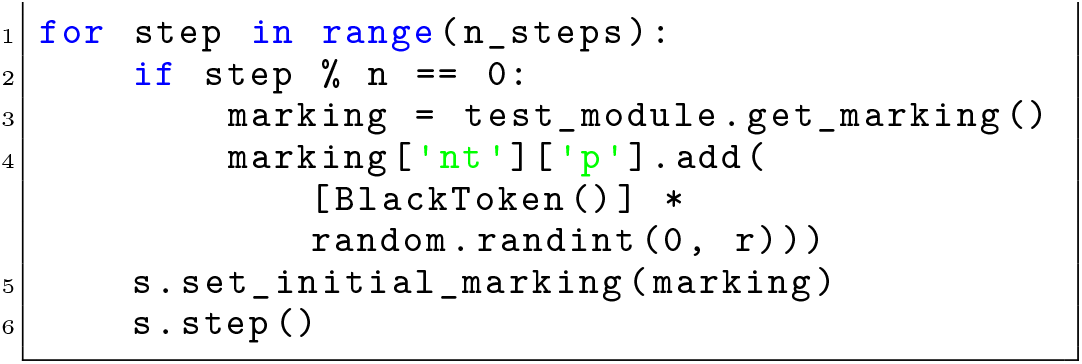
`nwn-petrisim` stimuli administration

In the stochastic simulation supported by `nwn-petrisim`, conflicts among transitions competing for tokens are managed through the randomized ordering of transitions enabling and firing events. All transitions have a user-defined firing probability of `p` set by default to 0.6. Furthermore, for each firing event, the set of tokens consumed by the transition is randomly chosen from those available in the input `place`. These random selections prevent the systematic exclusion of specific transitions from firing.

## IV. Results and Discussion

The BiSDL allows synthetic biologists to quickly model and design multicellular synthetic systems, simulating their behaviour. Synthetic biology aimed first to develop essential genetic constructs to control specific intracellular processes, then to combine such essential elements into complex circuits within or across cells [52]. Complex circuits enable a broader range of controllable behaviours yet have the drawback of metabolic burden and unknown interactions at the host. To address these constraints, synthetic biology has shifted focus towards designs based on multicellular networks [53], where splitting the overall construct across different cells facilitates integration into host cells and limits their metabolic burden. Construct parts interact *via* intercellular communication, and the desired behaviour emerges from the interaction between the different cells. Multicellular synthetic designs must consider complex interactions between the construct and the host cells. Results prove BiSDL capability for (1) model description and (2) exploration of system behaviour over three case studies of multicellular synthetic designs: a bacterial consortium (see subsection IV-A), a synthetic morphogen system (see subsection IV-B), and a conjugative plasmid transfer (see subsection IV-C).

### A. Case study 1 - bacterial consortium

The first case study focuses on implementing gene expression control across different bacterial cells. To achieve this, a synthetic biologist can exploit gene expression regulation across cells operated by the lactose repressor protein (LacI).

Initially, the biologist must identify a reliable source of knowledge regarding synthetic designs that realize the desired behaviour. The Registry for Standard Biological Parts holds a collection of predefined genetic constructs with known functionality [54]. These constructs can serve as templates for the DBTL process. Selecting and combining these parts makes it possible to design a bacterial consortium where the overall genetic device enforces LacI-operated gene expression regulation across cells. This consortium comprises two cell types. *Controller cells* establish baseline 3-oxohexanoyl-homoserine lactone (3OC6HSL) production, inhibited by a reference signal: LacI administration (Figure 4, top panel A). Conversely, *Target cells* initiate Green Fluorescent Protein (GFP) reporter signal production only when receiving the Acylhomoserine lactones (AHL) molecular signal, which, in this system, is 3OC6HSL (Figure 4, bottom panel B).

**Fig. 4.**
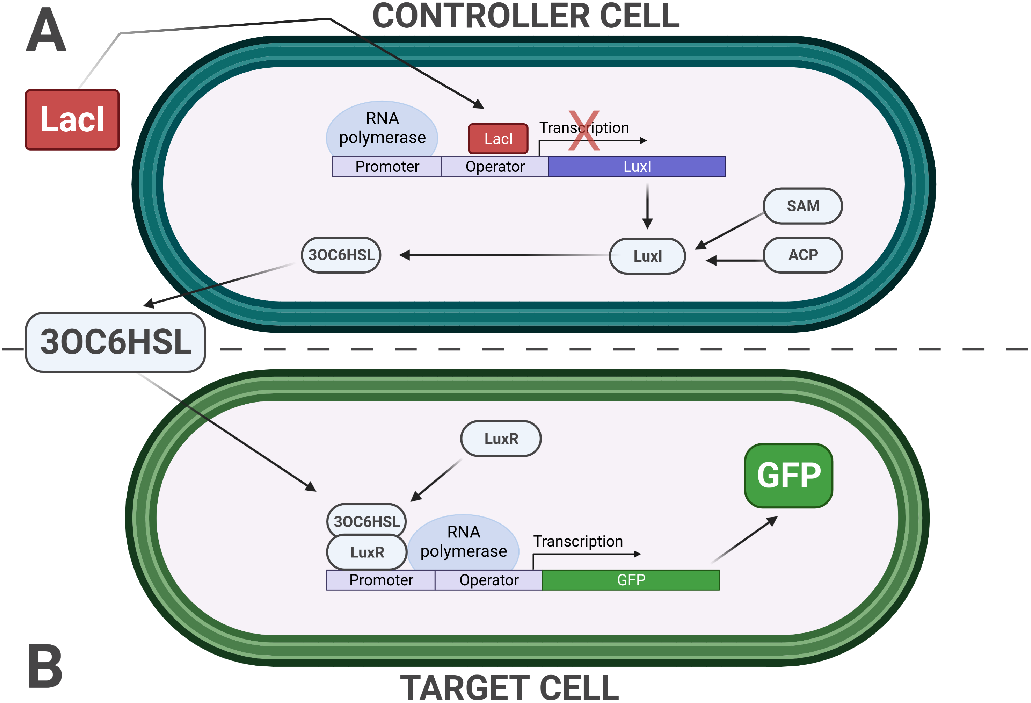
The multicellular bacterial consortium synthetic design was considered for the first case study. This consortium comprises two cell types. (A) *Controller cells* establish baseline 3OC6HSL production, inhibited by a reference signal (LacI administration); (B) *Target cells* initiate GFP reporter signal production only when receiving the AHL molecular signal, which, in this system, is 3OC6HSL.

Various Standard Biological Parts contribute to the design. *Part:BBa_C0012* (LacI protein) serves as the reference signal, inhibiting the Lac-repressible promoter. *Part:BBa_I13202* (3OC6HSL Sender Controlled by Lac Repressible Promoter) integrated with S-adenosylmethionine (SAM) and an acylated acyl carrier protein (ACP) substrate to synthesize 3OC6HSL [55] constructs in the *Controller cell. Part:BBa_E0040* (GFP) together with *Part:BBa_T9001* (Producer Controlled by 3OC6HSL Receiver Device) complete the design with the inducible reporter gene expression in the *Target cell*. The GFP levels serve as the readout signal.

The synthetic biologist who leverages BiSDL to describe the synthetic bacterial consortium should model the synthetic construct split across Controller and Target cells. Moreover, the model must include the biological interactions and mechanisms involved, such as transcriptional processes (gene expression), activation and inhibition of gene expression, protein production and degradation, enzymatic reactions, and inter-cellular signaling. algorithm 3 provides a BiSDL description of the synthetic bacterial consortium that considers all of these relevant aspects.

The illustrated bacterialConsortium `MODULE` contains two `SCOPE` statements: one for the Controller cell (`producer`) (algorithm 3, lines 3-20); another one for the Target cell (`sensor`) (algorithm 3, lines 21-34). Each `SCOPE` definition includes its name and two-dimensional coordinates on the spatial grid underlying the model (algorithm 3, lines 3 and 21). Each `SCOPE` contains a single `PROCESS` representing the fundamental biological functionality of each cell: `AHL_production` for the `producer` and `GFP_production` for the `sensor`. The `TIMESCALE` at the top level (algorithm 3, line 2) indicates the base pace for the `bacterialConsortium`. On the other hand, the `TIMESCALE` of each `PROCESS` (algorithm 3, lines 5 and 22) indicates the process slowdown factor related to the base pace: `AHL_production` evolves at half the base pace and `GFP_production` evolves at one-third of the base pace. The `bacterialConsortium` also contains declarations of the `DIFFUSION` processes that set up bidirectional connections between the two `SCOPE` constructs (`producer` and `sensor`) and the diffusion of `AHL_molecule` across them (algorithm 3, lines 33-37).

#### Algorithm 3: BiSDL description of the bacterial consortium.

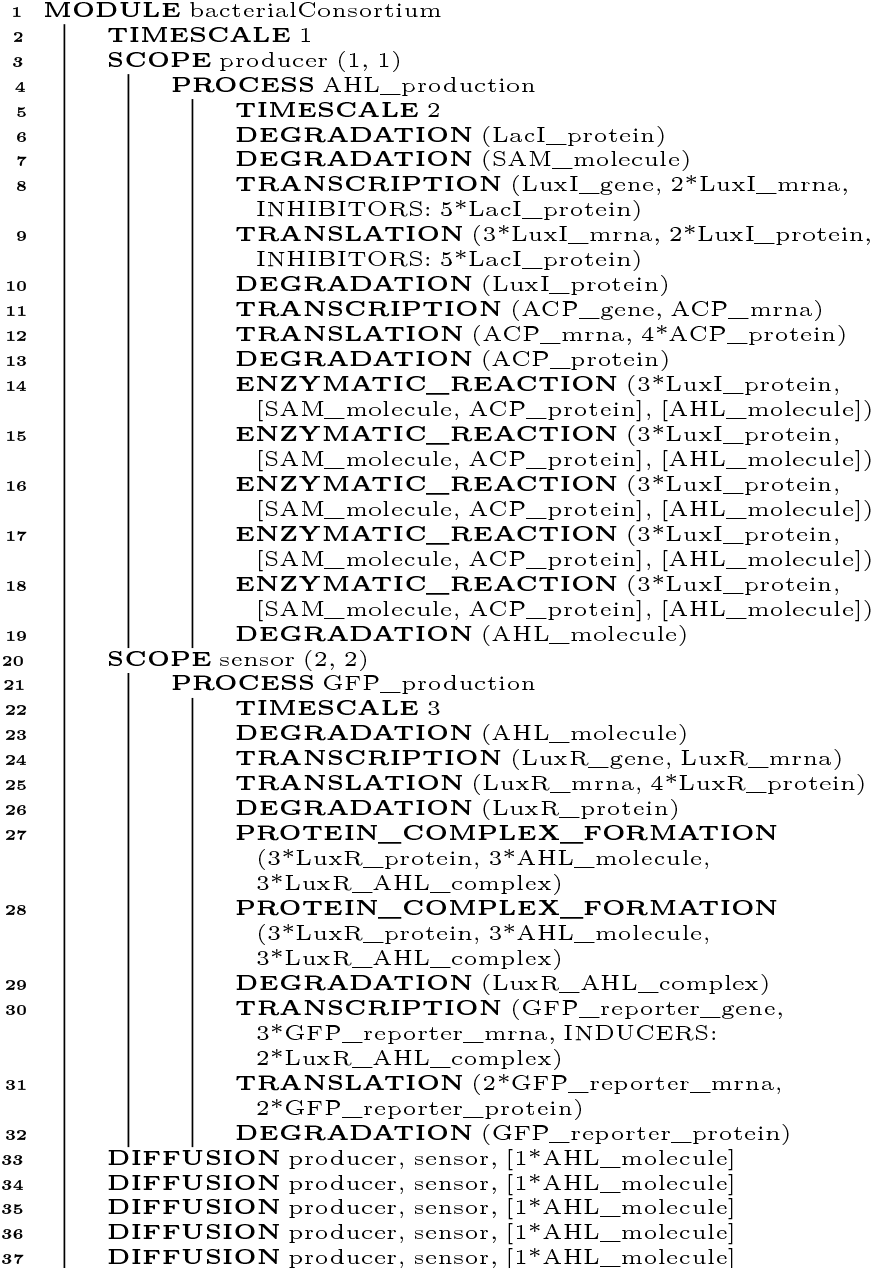

The BiSDL supports a very compact description of the system, using approximately 25% of the lines of code required by the low-level `nwn-snakes` model: 50 lines of code (see algorithm 3) versus 203 lines of code in the compiled `nwn-snakes` Python model file (based on the files in the public `GitHub` repository, see Availability of data and materials).

To compile the BiSDL description into a `nwn-snakes` model, the synthetic biologist uses the BiSDL compiler (see subsection III-D) to generate a `nwn-snakes` file that contains all the NWN models required by the BiSDL description. Visualization of the NWN models relies on the `GraphViz` visualization tool [56], provided by `SNAKES` as a plugin. For this use case, the NWN description includes a top-level net, where two places contain one net token each, and a bottom-level net where these net tokens lie.

Figure 5 visualizes the top-level NWN model, where the places that contain net tokens correspond to the two BiSDL `SCOPE` statements. The *producer* place holds the *AHL_production* net token (Figure 6), while the *sensor* place holds the *GFP_production* net token (Figure 7). Several transitions connect the two places, allowing the bidirectional diffusion of *AHL_molecule* colored tokens across them and the net tokens they contain during simulation, thanks to the `nwn-snakes` synchronization and communication capabilities (see subsubsection III-D1).

**Fig. 5.**
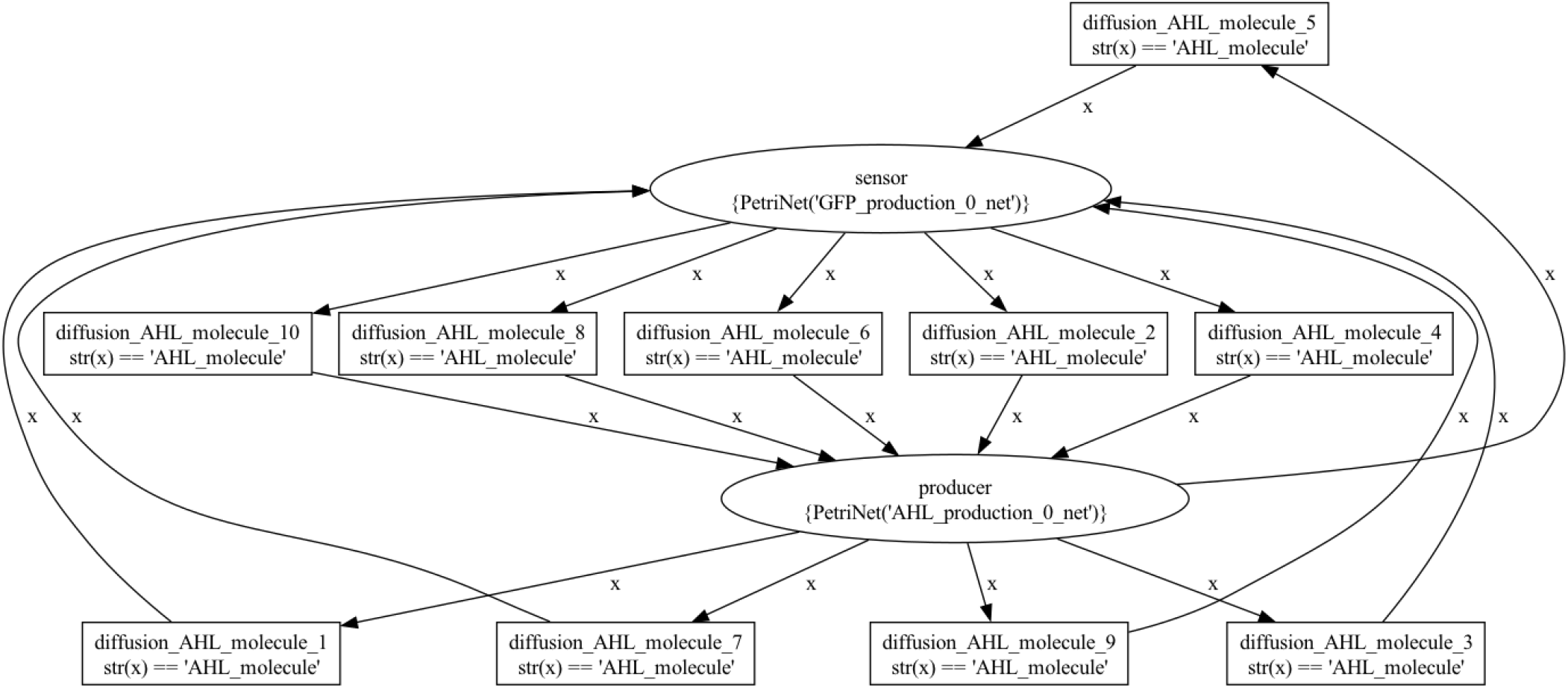
The top-level bacterial_consortium net architecture. The places that contain net tokens correspond to the two BiSDL `SCOPE` statements. The *producer* place holds the *AHL_production* net token, while the *sensor* place holds the *GFP_production* net token. Several transitions connect the two places, allowing the bidirectional diffusion of *AHL_molecule* colored tokens across them and the net tokens they contain during simulation, thanks to the `nwn-snakes` synchronization and communication capabilities.

**Fig. 6.**
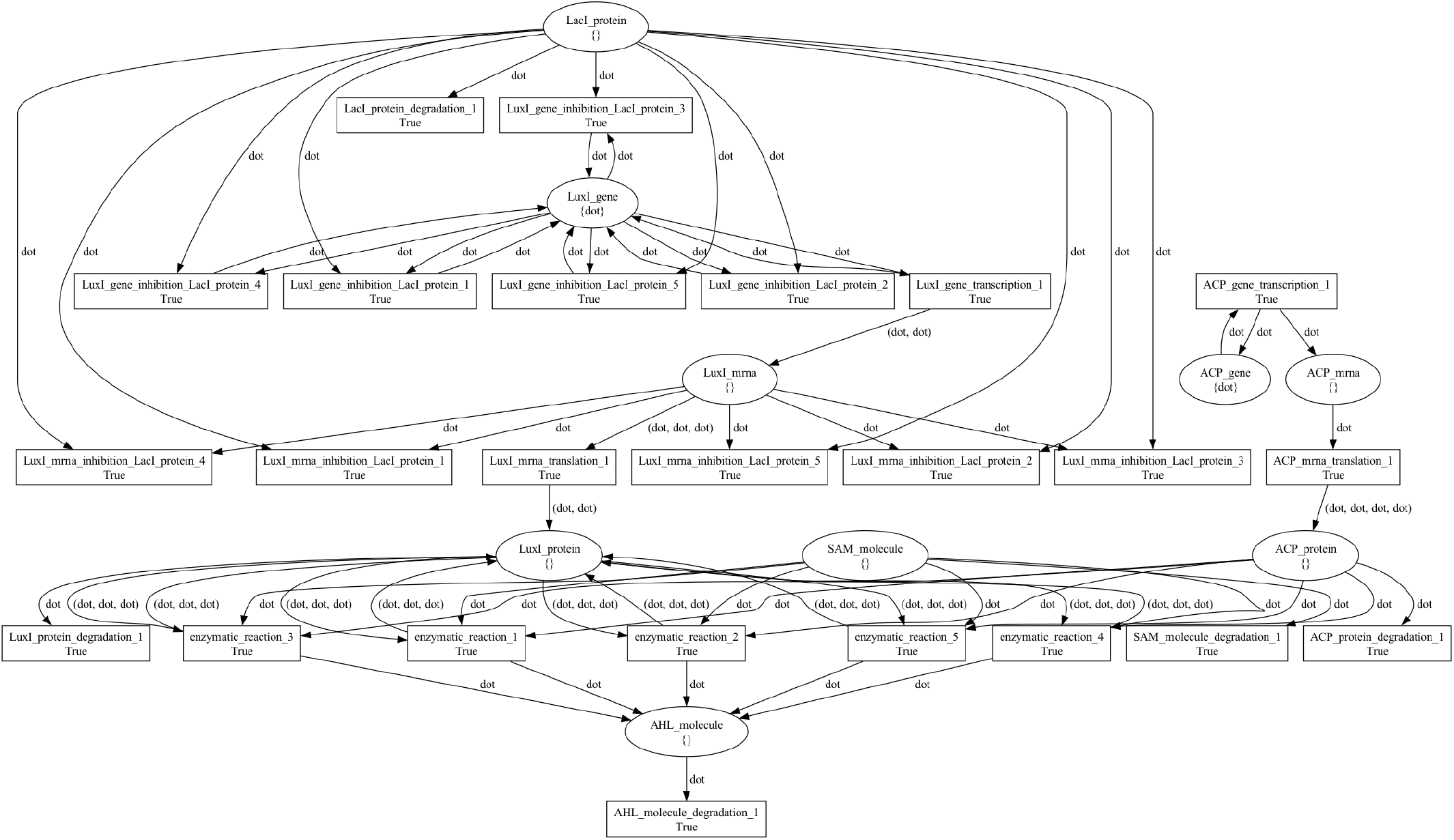
The producer bacterial_consortium net token architecture. Places model genes, transcripts, proteins, and molecules, while transitions model processes involving them, such as transcription, translation, degradation, and enzymatic reactions. Black tokens model discrete quantities of resources in each place and are represented by the `dot` symbol.

**Fig. 7.**
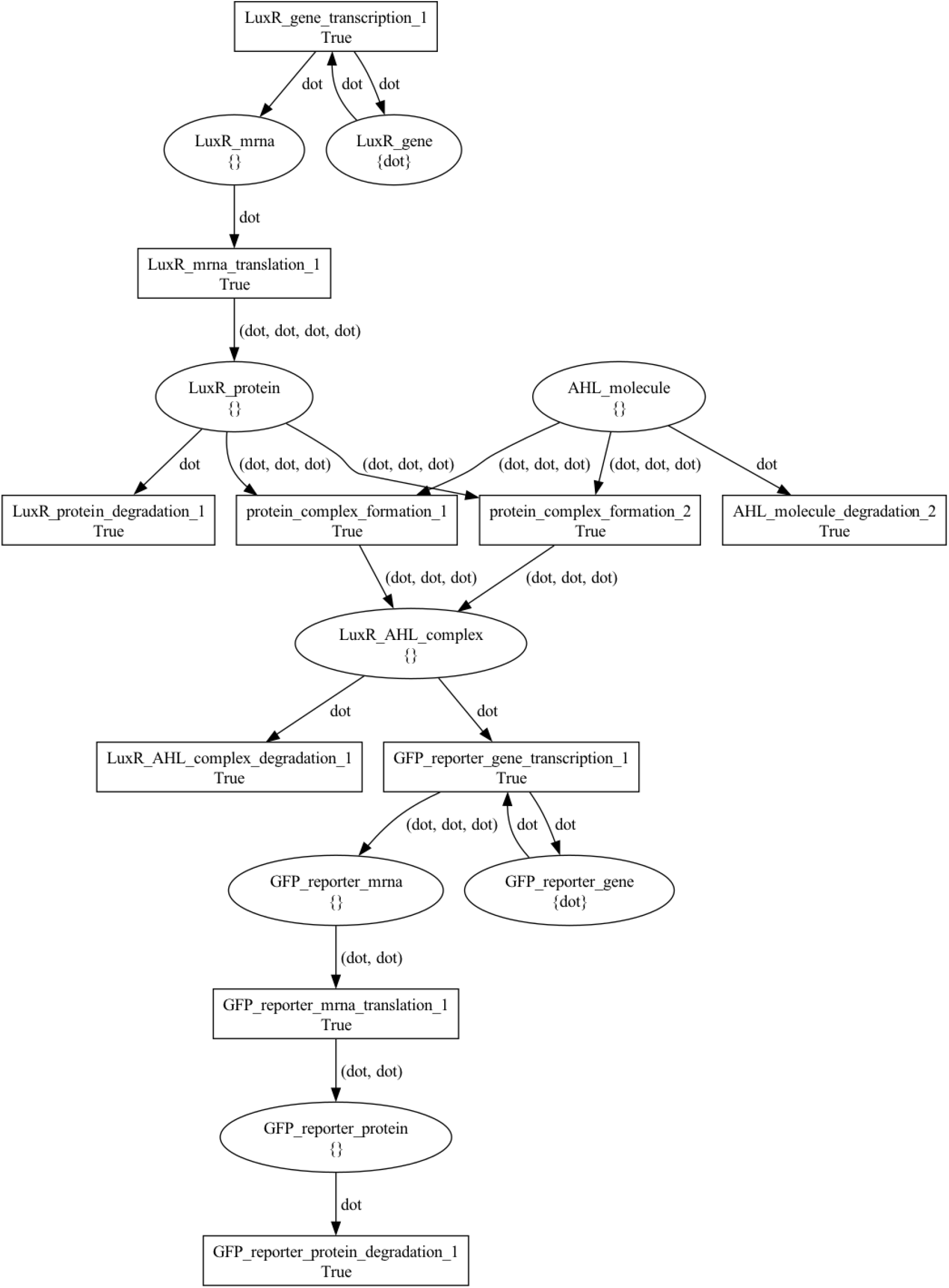
The sensor bacterial_consortium net token architecture. Places model genes, transcripts, proteins, and molecules, while transitions model processes involving them, such as transcription, translation, degradation, and enzymatic reactions. Black tokens model discrete quantities of resources in each place and are represented by the `dot` symbol.

Figure 6 and Figure 7 visualize the *producer* and *sensor* net tokens, respectively. In these PN, places model genes, transcripts, proteins, and molecules, while transitions model processes involving them, such as transcription, translation, degradation, and enzymatic reactions. Black tokens model discrete quantities of resources in each place and are represented by the `dot` symbol. The simulation of BiSDL-compiled `nwn-snakes` models shows that LacI levels control *GFP_protein* levels, consistently with the expected behaviour under the following conditions:

- *noLacI* : the absence of LacI administration;
- *lowLacI* : constant and low LacI administration (n=3 and r=3);
- *highLacI* : constant and high LacI administration (n=3 and r=10);

Values of `n` and `r` determine the intensity of stimulus administration (see subsubsection III-D2).

Figure 8 presents the marking evolution of the *LacI* signal mediators (`Lux1_protein` and `AHL_molecule`) and Target (`GFP_reporter_protein`) in the bacterial consortium after the three considered LacI administration schemes. *noLacI* (Figure 8, top left panel A) does not interfere with `AHL_molecule` levels (Figure 8, middle left panel D), inducing transcription of high `GFP_reporter_protein` readout signals (Figure 8, bottom left panel G). *lowLacI* (Figure 8, top central panel B) hampers `AHL_molecule` levels (Figure 8, middle central panel E), resulting in the transcription of lower `GFP_reporter_protein` (Figure 8, bottom central panel H). *highLacI* (Figure 8, top right panel C) almost shuts down `AHL_molecule` levels (Figure 8, middle right panel F), suppressing the transcription of high `GFP_reporter_protein` readout signals almost completely (Figure 8, bottom right panel I). The results show consistency with the expected system behavior, an inverse relation between LacI stimulus and readout signal levels.

**Fig. 8.**
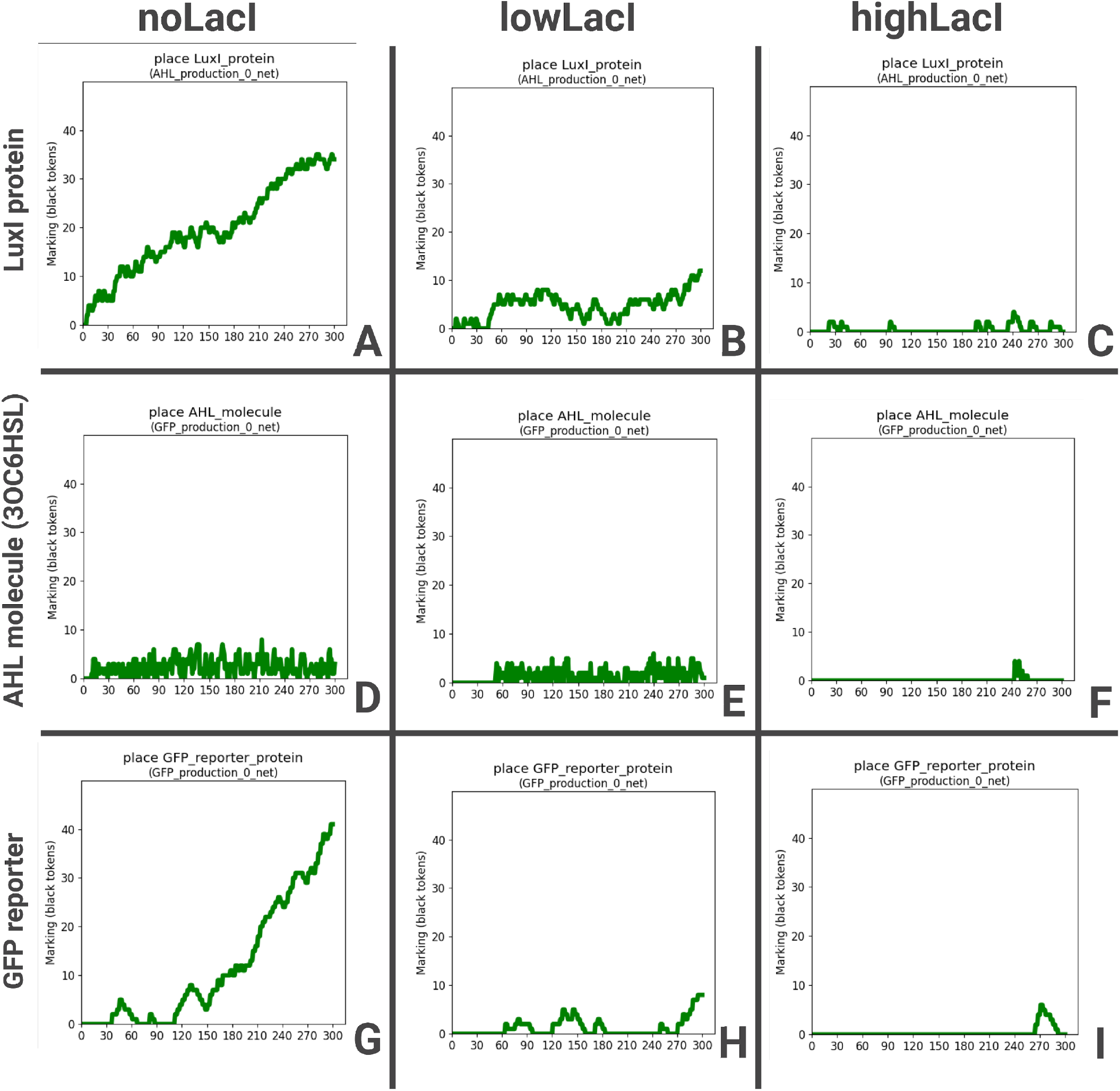
Marking evolution of the `LacI` signal mediators (`Lux1_protein and AHL_molecule`) and Target (`GFP_reporter_protein`) in the bacterial consortium after three LacI administration schemes. (A) *noLacI* does not interfere with (D) `AHL_molecule` levels, (G) inducing transcription of high `GFP_reporter_protein` readout signals. (B) *lowLacI* (E) hampers `AHL_molecule` levels, resulting in (H) the transcription of lower `GFP_reporter_protein`. (C) *highLacI* (F) almost shuts down `AHL_molecule` levels, (I) suppressing the transcription of high `GFP_reporter_protein` readout signals almost completely. The results show consistency with the expected system behavior, an inverse relation between LacI stimulus and readout signal levels. size, but they are available from the

### B. Case study 2 - RGB synthetic morphogen system

The second case study implements a synthetic morphogen system where the spatial interactions and organization of the cells sustain the emergence of a spatial pattern of red, green, and blue (RGB) fluorescent markers. In developmental processes, morphogens transmit positional signals to cells, diffusing from a source to create concentration gradients. Cells interpret these gradients using diverse signaling mechanisms, including paracrine signaling (short-range) and juxtacrine communication (cell-to-cell). Synthetic biology offers the potential to manipulate these mechanisms, enabling control over spatial arrangement and functional features in synthetic morphogenetic designs. This second case study illustrates how BiSDL descriptions express the essential dynamics underlying a multicellular synthetic design accounting for the role of spatial organization and neighborhood relations among cells.

This time, rather than relying on Standard Parts (refer to subsection IV-A), the objective is to replicate designs found in the scientific literature, such as the modular synNotch system outlined in [57], providing a platform for engineering orthogonal juxtacrine signaling, which functions independently of natural cellular communication pathways. It enables specific and controlled cell interactions, featuring an extracellular recognition domain, the Notch core regulatory domain, and an intracellular transcriptional domain. With the incorporation of fluorescent markers, the synNotch system proves to be a valuable tool for engineering multicellular synthetic systems.

This second case study comprises a three-layer multicellular circuit in which the Receiver cells (Cells B, Figure 9, top right panel B) inducibly expresses E-cadherin (*Ecad*_*hi*_) and a modified form of GFP working as a synNotch ligand on the cell membrane (*GFP*_*lig*_). Furthermore, *GFP*_*lig*_ serves as both a fluorescent reporter and a ligand for a secondary synNotch receptor with the cognate anti-GFP binding domain (Figure 9, central right panel C). The Sender cells (Cells A, Figure 9, top left panel A) constitutively express BFP, CD19 ligand, and the anti-GFP synNotch receptor, which drives expression of a low amount of E-cadherin (*Ecad*_*lo*_) fused with a mCherry reporter for visualization (Figure 9, central left panel D). Thus, Cells A have low adherence, blue fluorescence, and inducible red fluorescence (Figure 9, bottom left panel E), while Cells B have inducible green fluorescence and high adherence (Figure 9, bottom right panel F).

**Fig. 9.**
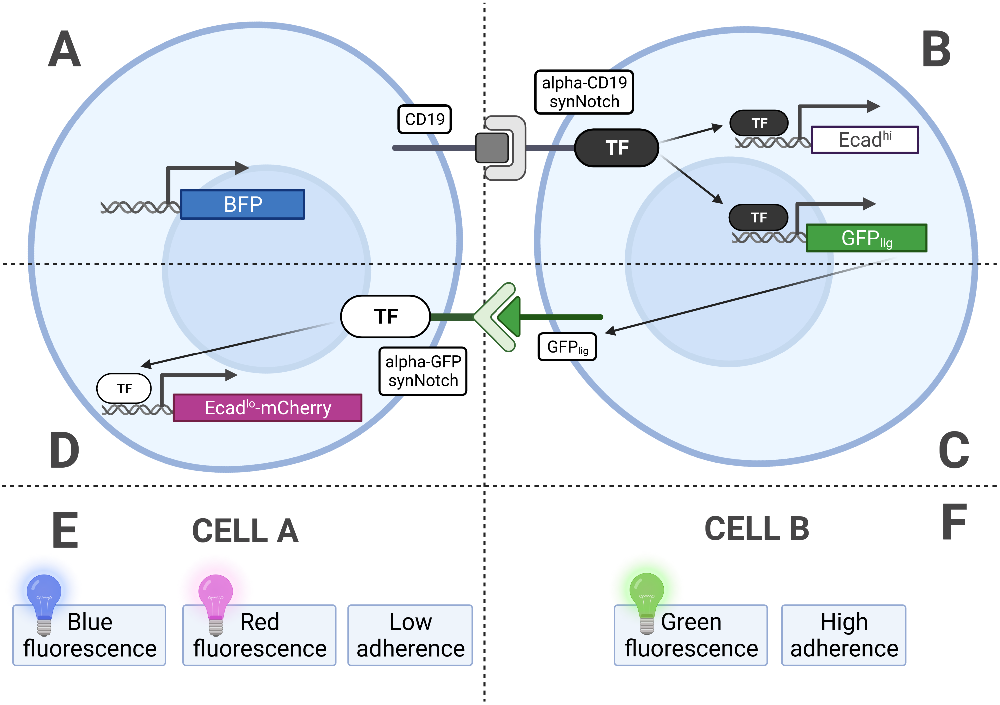
The multicellular synthetic circuit mediated by cell-cell communication makes the RGB pattern emerge. (A) The Sender cells (Cells A) constitutively express BFP, CD19 ligand, and the anti-GFP synNotch receptor, which drives expression of a low amount of E-cadherin (*Ecad*_*lo*_) fused with a (D) mCherry reporter for visualization. (B) Receiver cells (Cells B) inducibly express E-cadherin (*Ecad*_*hi*_) and a modified form of GFP working as a synNotch fluorescent reporter and a ligand for a secondary synNotch receptor with the cognate anti-GFP binding domain. (E) Cells A have low adherence, blue fluorescence, and inducible red fluorescence; (F) Cells B have inducible green fluorescence and high adherence. Adapted from [57].

Cells start as a disorganized aggregate (Figure 10, bottom left panel A), and CD19 in Cells A activates anti-CD19 synNotch in Cells B, inducing the expression of a high level of E-cadherin (*Ecad*_*hi*_) and *GFP*_*lig*_ (Figure 10, top left panel B). Cells B thus aggregate and form a compact group in the middle of the aggregate (Figure 10, bottom central panel C). The *GFP*_*lig*_ on Cells B activates the anti-GFP synNotch receptors on Cells A in direct contact with the central group, inducing *Ecad*_*lo*_ and the mCherry reporter (Figure 10, top right panel D), and making a spatially organized pattern of cells emerge in a synthetic morphogenetic pattern (Figure 10, bottom right panel E) with three concentric layers: a green internal core (Cells B expressing *Ecad*_*hi*_ and *GFP*_*lig*_) with high cell-cell adhesion, an outer layer of blue cells (Cells A expressing BFP), and a population of red cells in the middle layer (Cells A expressing *Ecad*_*lo*_ and mCherry).

**Fig. 10.**
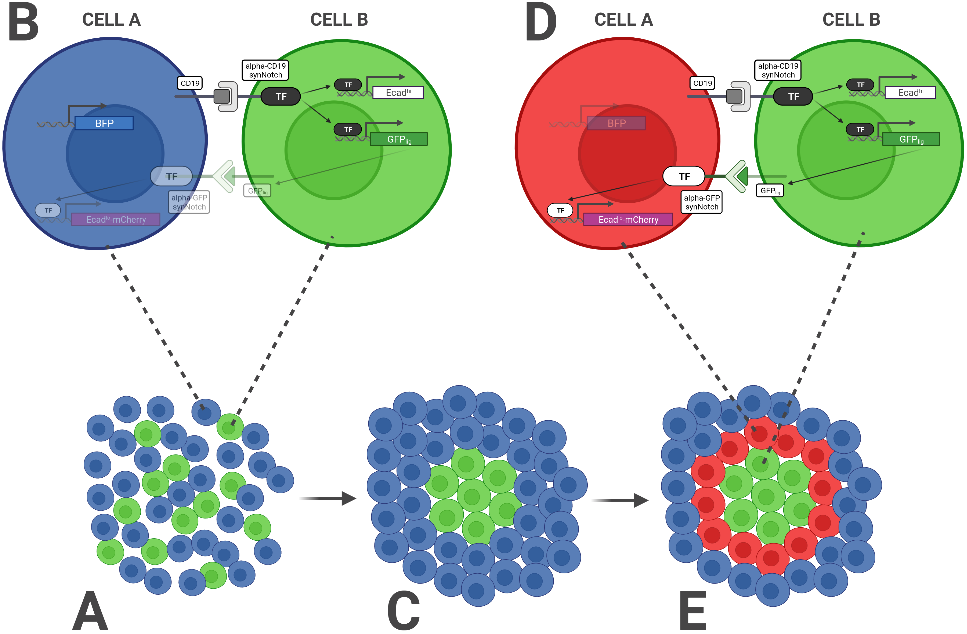
The RGB synthetic morphogenesis process for the second case study. (A) Cells start as a disorganized aggregate; (B) CD19 in Cells A activates anti-CD19 synNotch in Cells B, inducing the expression of a high level of E-cadherin (*Ecad*_*hi*_) and *GFP*_*lig*_; (C) Cells B aggregate and form a compact group in the middle of the aggregate; (D) The *GFP*_*lig*_ on Cells B activates the anti-GFP synNotch receptors on Cells A in direct contact with the central group, inducing *Ecad*_*lo*_ and the mCherry reporter, (E) making a spatially organized pattern of cells emerge in a synthetic morphogenetic pattern with three concentric layers: a green internal core (Cells B expressing *Ecad*_*hi*_ and *GFP*_*lig*_) with high cell-cell adhesion, an outer layer of blue cells (Cells A expressing BFP), and a population of red cells in the middle layer (Cells A expressing *Ecad*_*lo*_ and mCherry).Adapted from [57].

algorithm 4 illustrates the BiSDL description of the RGB synthetic morphogen system, where a central cell induces patterning in its neighbors of the first and second-degree. Vertical dots indicate sections describing red and blue cells `SCOPE`s with identical structure as the ones presented but different spatial coordinates. The complete description and the visualization of the compiled `nwn-snakes` PN models are not included due to their size, but they are available from the BiSDL `GitHub` public repository (see Availability of data and materials).

BiSDL again supports a compact description of the system composed of 165 lines of code, approximately 9% of the corresponding compiled `nwn-snakes` model file, having 1807 lines of code (based on the files in the public `GitHub` repository, see Availability of data and materials).

Results recapitulate the emergence of the expected simplistic version of one of the synthetic morphogenetic patterns presented in [57]. Figure 11 depicts the evolving intensity of the fluorescent markers in each cell during a simulation of the `nwn-snakes` Python models compiled from the BiSDL description (see algorithm 4). At first (t = 10), the central cell slightly affects only one of its neighbors of the first degree, while other cells keep producing the BFP signal (Figure 11, top left panel A); the central cell engages all of its neighbors of the first degree, inducing the expression of mCherry in them, whose intensity evolves throughout the simulation (from t=20 to t=50, Figure 11, top central panel B, top right panel C, bottom left panel D, bottom central panel E); at t=60 all first-degree neighbors of the central cell express high levels of mCherry (Figure 11, bottom right panel F). On the contrary, the simulated deletion of *GFP*_*lig*_ results in a stable pattern with a central, colorless cell of type B and all its neighbors of the first and second degree constitutively expressing BFP (data not shown).

**Fig. 11.**
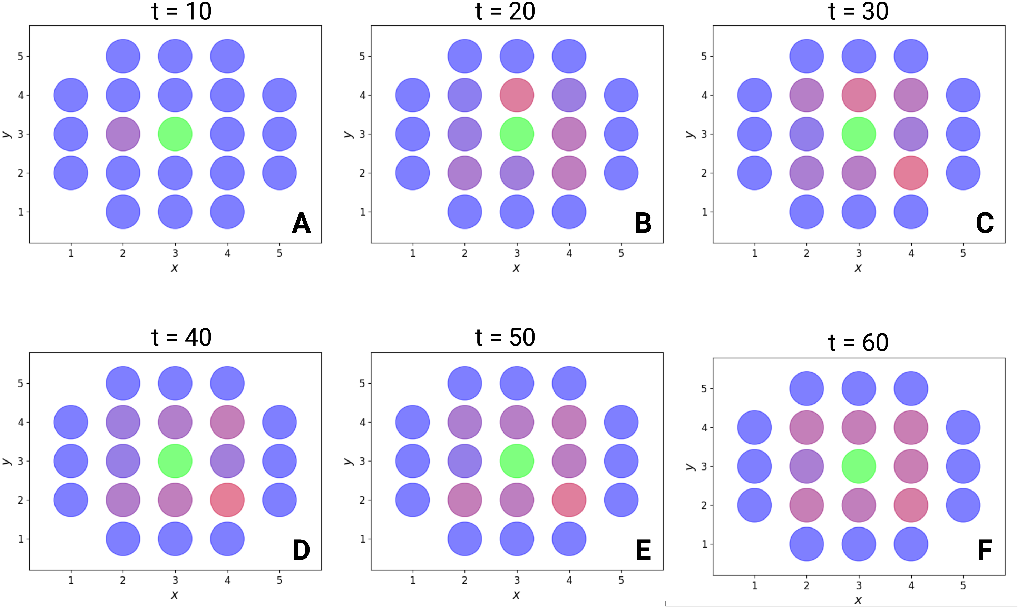
Evolution of the three fluorescent marker levels (GFP, BFP and mCherry) in each cell on the two-dimensional spatial grid along the simulation of the `nwn-snakes` Python models compiled from the BiSDL description. (A) At first (t = 10), the central cell slightly affects only one of its neighbors of the first degree, while other cells keep producing the BFP signal; (B-E) the central cell engages all of its neighbors of the first degree, inducing the expression of mCherry in them, whose intensity evolves throughout the simulation (from t=20 to t=50); (F) all first-degree neighbors of the central cell express high levels of mCherry (t=60).

#### Algorithm 4: BiSDL description of the RGB synthetic morphogen system.

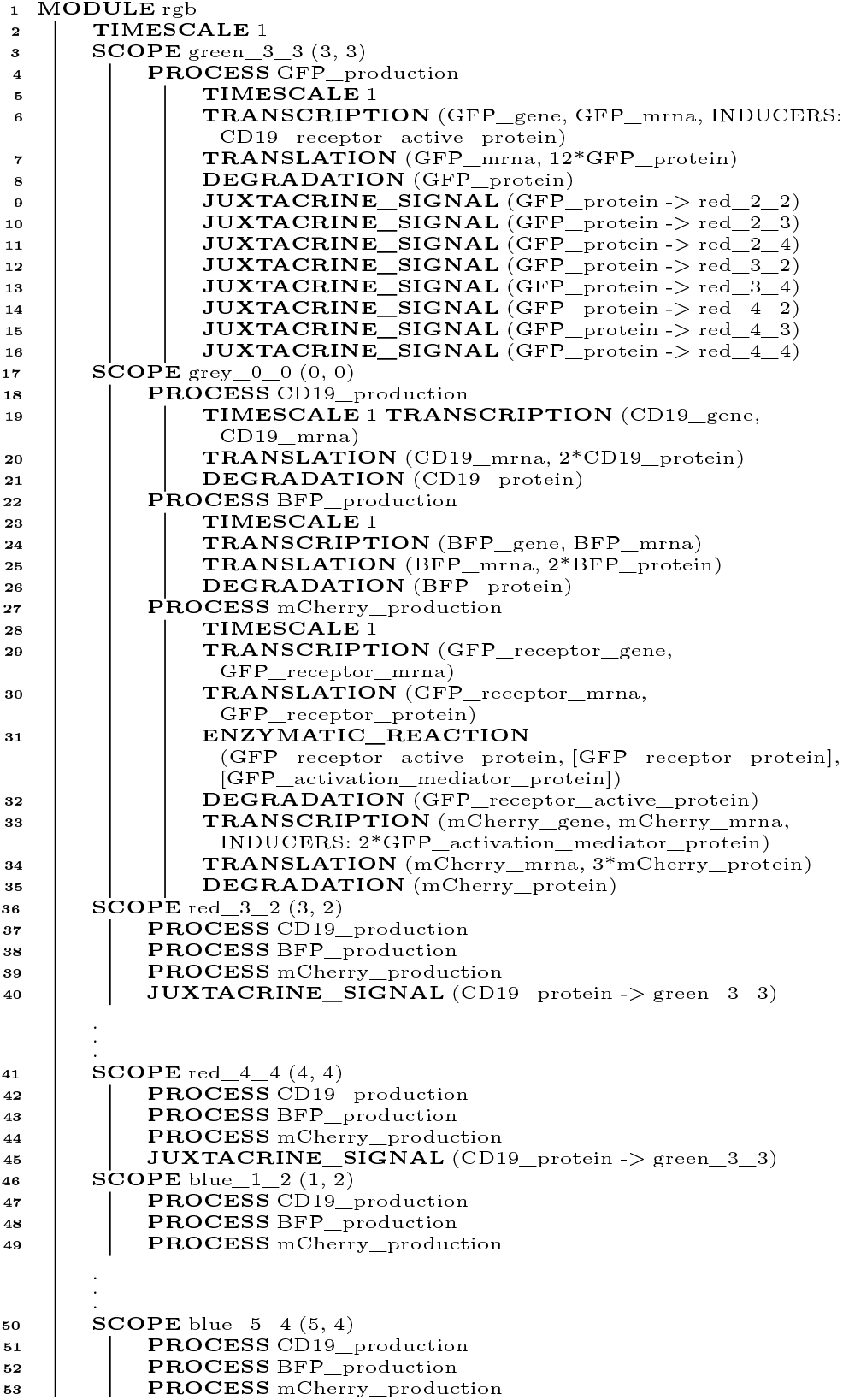

### C. Case study 3 - conjugative plasmid transfer

Plasmids are crucial in disseminating antibiotic resistance, virulence genes, and various adaptive traits within bacterial communities through horizontal gene transfer [61]. The third case study examines antibiotic resistance (R) conjugative plasmids transfer between bacterial cells, mirroring the fundamental conjugation mechanism presented in [58]. As depicted in Figure 12, the plasmid transfer process [59] initiates with a Donor cell harboring a conjugative R plasmid (Figure 12, top left). The Donor extends a Pilus, a proteinaceous protrusion [60], encoded by the R plasmid, to establish contact with a compatible recipient cell, referred to as the Transconjugant cell [58] (Figure 12, top right). Upon contact, the Pilus retracts, bringing the cells into proximity and forming a conjugation bridge. This bridge facilitates the transfer of one of the plasmid DNA strands in a linearized form from the Donor to the Transconjugant (Figure 12, center). Subsequently, both cells harbor a single-stranded copy of the R plasmid. Finally, through circularization and DNA synthesis, the Donor and Transconjugant cells complete the second strand for their respective R plasmid copies. Consequently, both cells become capable of further disseminating the R plasmid, effectively functioning as Donor cells (Figure 12, bottom), thereby facilitating the propagation of antibiotic resistance.

**Fig. 12.**
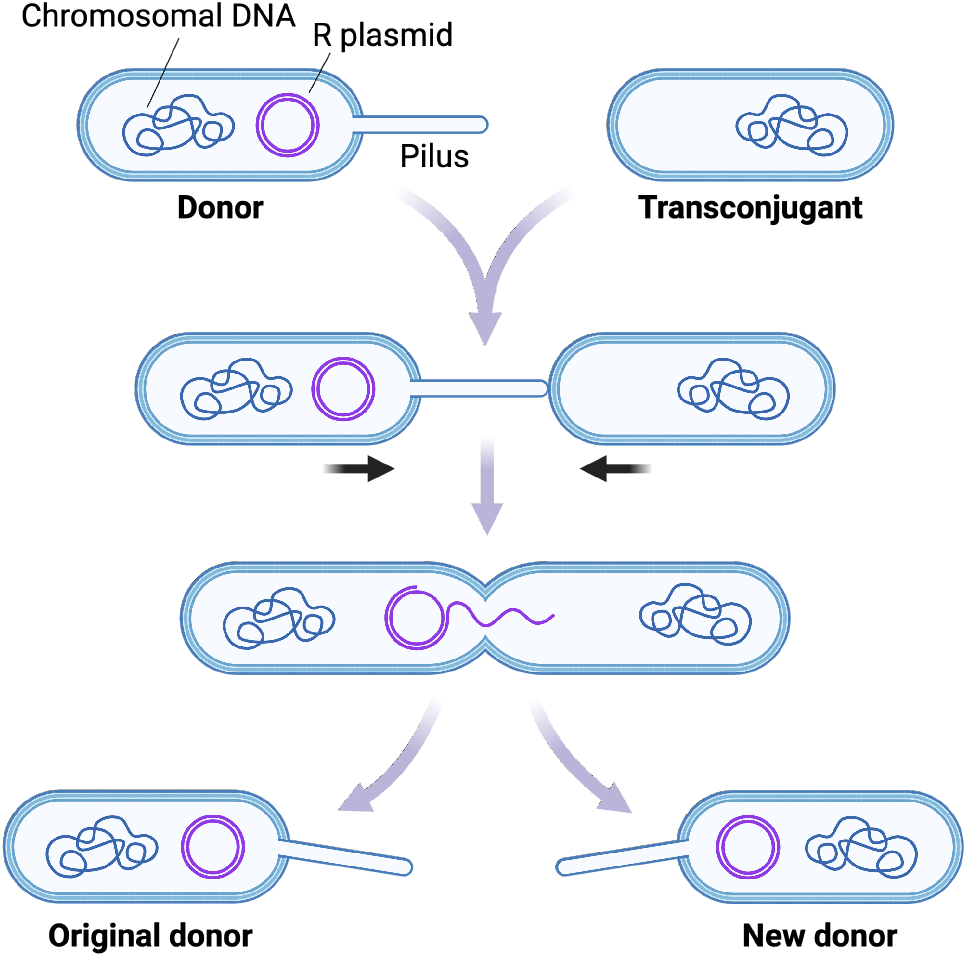
The mechanism considered for the third case study is antibiotic resistance (R) plasmid transfer across bacterial cells, recapitulating the basic conjugation mechanism modeled in [58]. The plasmid transfer mechanism [59] begins with a Donor cell that carries a conjugative R plasmid (top left). The Donor extends a Pilus, a proteinaceous protrusion [60] to contact a compatible receiver cell, named the Transconjugant cell (top right). In this example, the Pilus is encoded in the R plasmid. Upon contact, the Pilus retracts, pulling the cells together and establishing a conjugation bridge. This bridge enables the transfer of one of the plasmid DNA strands in linearized form from the Donor to the Transconjugant (center). At this point, both cells hold a single-stranded copy of the R plasmid. Finally, both cells build the second strand for their respective R plasmid copies through circularization and DNA synthesis. Thus, both Donor and Transconjugant cells are equipped to disseminate the R plasmid further, making them both Donor cells (bottom) enacting antibiotic resistance propagation.

algorithm 5 provides a BiSDL description of the conjugative R plasmid transfer from the Donor to the Transconjugant cell.

The illustrated plasmidTransfer `MODULE` contains three `SCOPE` statements: one for the Donor cell (`donor`, (algorithm 5, lines 3-15); another one for the Transconjugant cell (`transconjugant`, algorithm 5, lines 21-32), and a third one for the Pilus proteinaceous structure mediating the conjugation process (`pilus`, algorithm 5, lines 16-20). This shows the flexibility of the `SCOPE` construct in supporting the modeler for expressing different types of biological compartments. `JUXTACRINE_SIGNAL` mechanisms connect the `SCOPE` statements from the `donor` to the `transconjugant` through the `pilus`, recapitulating the R plasmid transfer process.

Figure 13 visualizes the top-level NWN model for this use case, having a place for each of the three BiSDL `SCOPE` statements, holding the *donor* net token (Figure 15), the *transconjugant* net token (Figure 16) and the *pilus* net token (Figure 14), respectively.

**Fig. 13.**
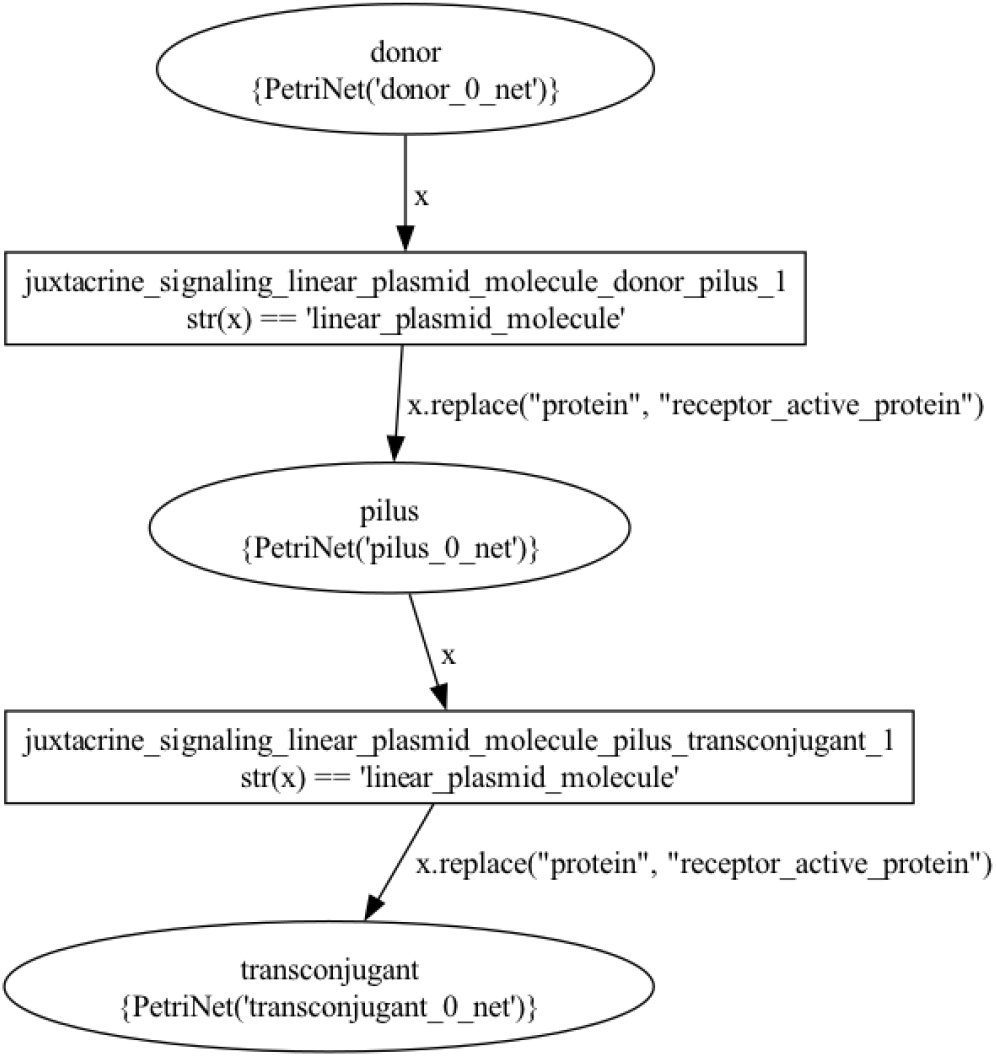
The top-level plasmid_transfer net architecture. The places that contain net tokens correspond to the two BiSDL `SCOPE` statements, holding the *donor* net token (Figure 15), the *transconjugant* net token (Figure 16) and the *pilus* net token (Figure 14), respectively.

**Fig. 14.**
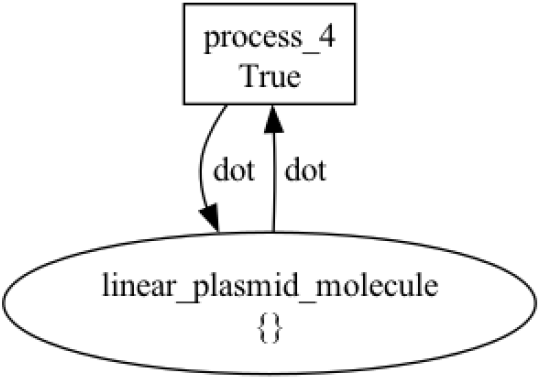
The *Pilus* plasmid_transfer net token architecture. Places model genes, transcripts, proteins, and molecules, while transitions model processes involving them, such as transcription, translation, degradation, and enzymatic reactions. The black tokens model discrete quantities of resources in each place and are represented by the `dot` symbol.

**Fig. 15.**
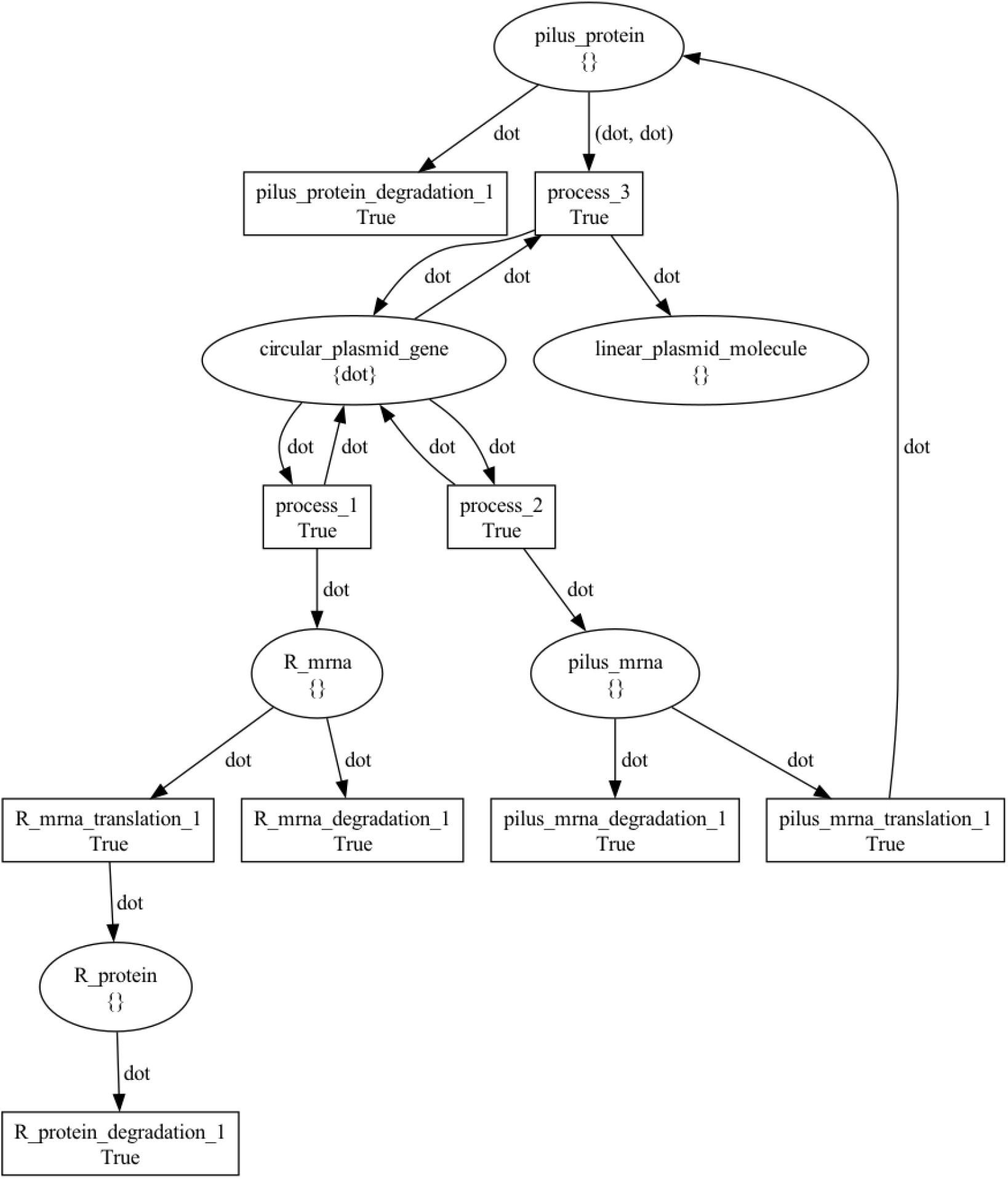
The *donor* plasmid_transfer net token architecture. Places model genes, transcripts, proteins, and molecules, while transitions model processes involving them, such as transcription, translation, degradation, and enzymatic reactions. Black tokens model discrete quantities of resources in each place and are represented by the `dot` symbol.

**Fig. 16.**
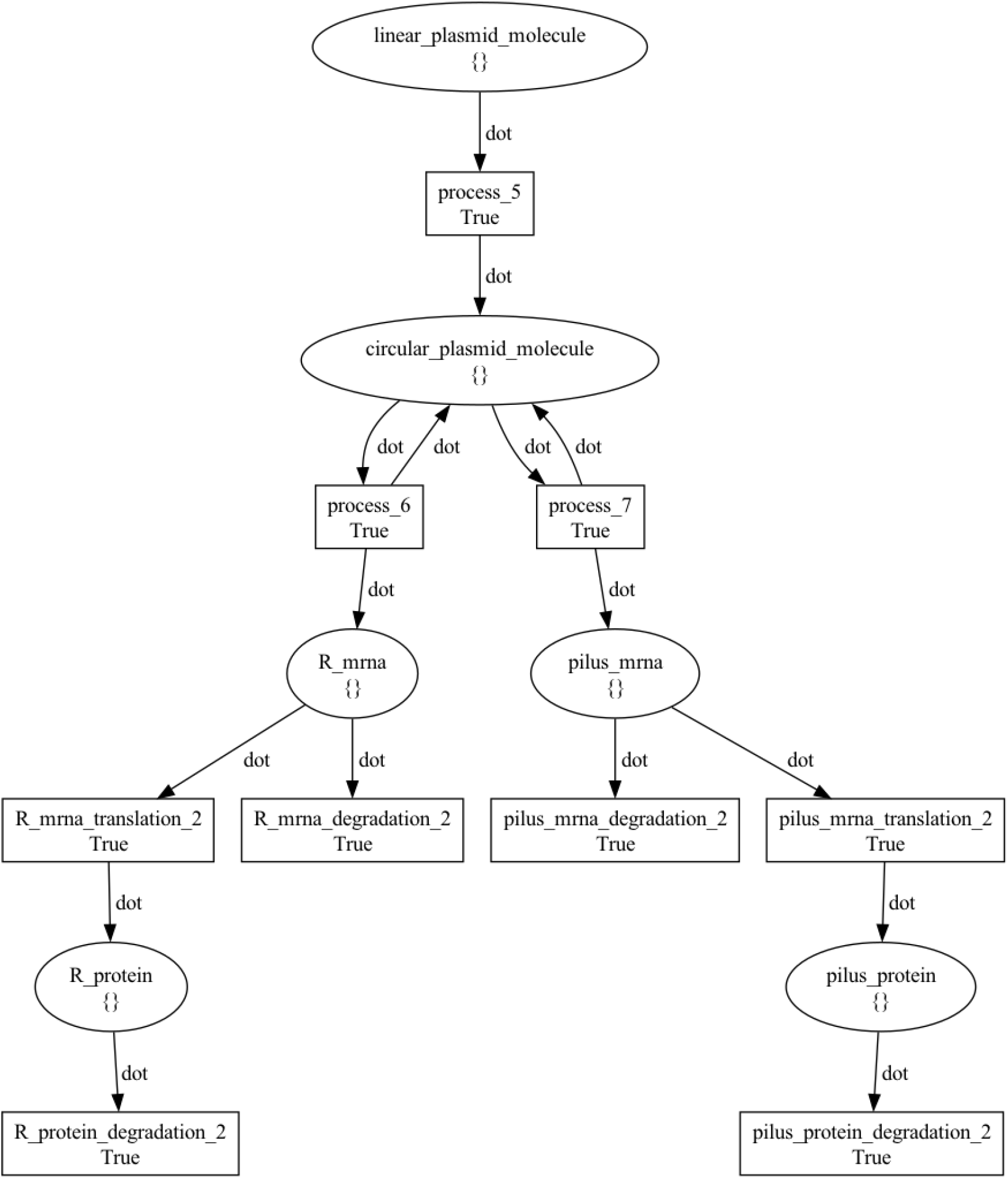
The *transconjugant* plasmid_transfer net token architecture. Places model genes, transcripts, proteins, and molecules, while transitions model processes involving them, such as transcription, translation, degradation, and enzymatic reactions. Black tokens model discrete quantities of resources in each place and are represented by the `dot` symbol.

The simulation of BiSDL-compiled `nwn-snakes` models demonstrates that transferring R plasmids from the Donor to the Transconjugant cells aligns with anticipated behavior. Throughout the simulation, the Donor cell consistently retains one R plasmid (Figure 17, top left), while the Transconjugant cell initially lacks any R plasmids (Figure 17, top right). The R plasmid within the Donor encodes the Pilus protein, which initiates the conjugation process upon translation. The Pilus mediates two R plasmid transfer events, illustrated by linearized single-strand R plasmids within it (Figure 17, center), resulting in two subsequent increases in R plasmid copies within the Donor cell (Figure 17, top right). R plasmids encode the R protein, conferring antibiotic resistance. The R protein is consistently present in the Donor cell due to the R plasmid (Figure 17, bottom left). Conversely, in the Transconjugant cell, R protein levels only rise above zero after the initial R plasmid transfer event (Figure 17, bottom right). These findings indicate that the simulation accurately reproduces the conjugative plasmid transfer process, leading to R protein-mediated antibiotic resistance propagation.

**Fig. 17.**
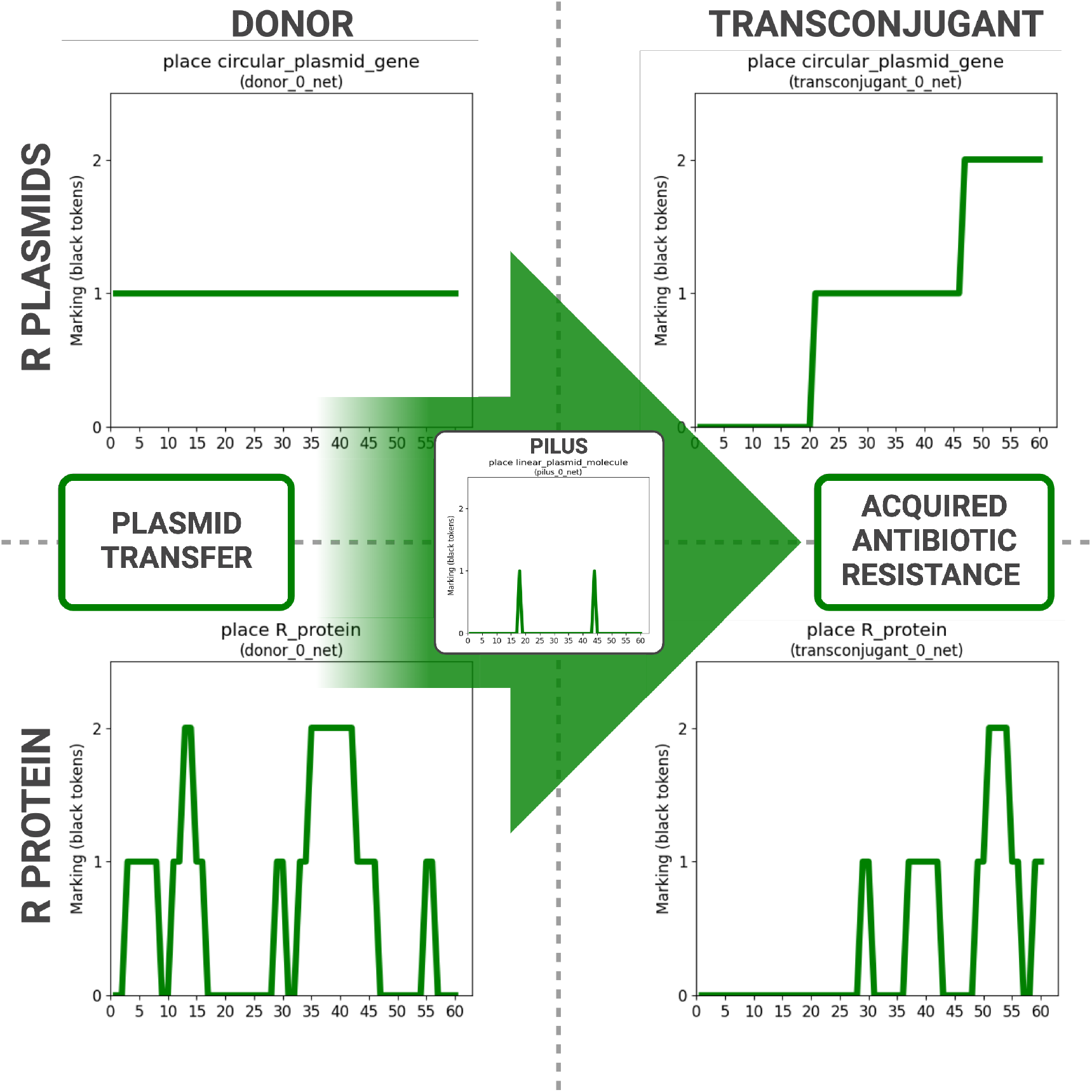
Simulation of BiSDL-compiled `nwn-snakes` models shows that the transfer of R plasmids from the Donor to the Transconjugant cells is consistent with the expected behavior. The Donor cell holds one R plasmid throughout the simulation (top left), while the Transconjugant cell starts with none (top right). The R plasmid in the Donor encodes for the Pilus protein, which initiates the conjugation process as soon as it is translated. The Pilus mediates two R plasmid transfer events, depicted as the presence of a linearized single-strand R plasmid within it (center), resulting in two subsequent increases of R plasmid copies in the Donor cell (top right). R plasmids encode for R protein, which provides antibiotic resistance. In the Donor cell, the R protein is present from the start due to the presence of the R plasmid (bottom left). Otherwise, protein levels rise above zero in the Transconjugant cell R only after the first R plasmid transfer event (bottom right). These results show that simulation recapitulates the plasmid-transfer conjugation process, causing the acquisition of R protein-mediated antibiotic resistance.

#### Algorithm 5: BiSDL description of the conjugative R plasmid transfer from the Donor to the Transconjugant cell.

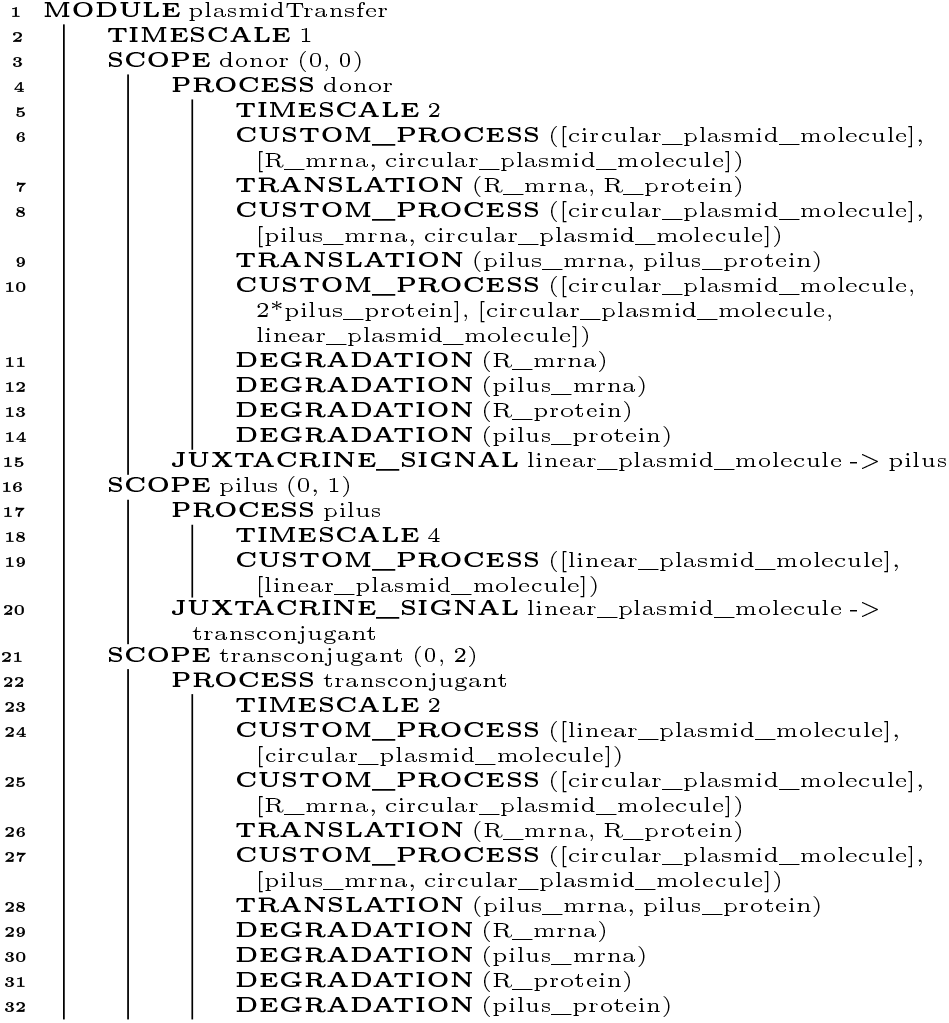

## V. Conclusions

In conclusion, the BiSDL framework represents a significant advancement in synthetic biology modeling. The core aim of BiSDL is to merge the detailed expressive capabilities of computational models with the user-friendliness of high-level languages, providing a tool that is both powerful and more accessible than a low-level language. In fact, BiSDL requires only basic modeling and programming skills compared to the advanced modeling and programming skills required by low-level languages. The hierarchical and modular structure of BiSDL is ideal for capturing the inherent complexity of biological systems and constructing reusable and adaptable components.

Through the development of a prototype BiSDL compiler, models can be compiled into low-level descriptions that encapsulate the spatial, hierarchical, and dynamic behaviors of biological entities, using the NWN approach to handle biological complexity. As BiSDL closely mirrors the domain-specific language used within the biological domain, the compiler closes the gap between high-level biological semantics and NWN low-level formalism syntax. Generated models rely on the `nwn-snakes` library, an extension of the `SNAKES` library [51] to support the NWN formalism. Results indicate that BiSDL can dramatically simplify complex model descriptions, significantly reducing the code needed to represent sophisticated systems. This is evidenced by the bacterial consortium, RGB and conjugative plasmid transfer case studies.

The accompanying `nwn-petrisim` simulator has been developed to reproduce and investigate behaviors of systems modeled in BiSDL, confirming that the BiSDL-compiled models accurately represent the expected behaviors of the systems studied.

Future developments may integrate into BiSDL a tool for formal verification [62], inherently supported by PN as a low-level formalism. This would allow the validation of complex models by rigorously checking for correctness according to specific criteria. Several integrable tools exist for PN formal verification, including TINA (Time Petri Net Analyzer) [63], ABCD plus Neco for SNAKES models [64] and GreatSPN [65], which provides a user-friendly visual interface, in alignment with BiSDL’s aim for broader accessibility.

The proposed BiSDL constructs exemplify the expressiveness of the language, and its closeness to the biological semantics. Future developments include gradually extending the syntax of the language with additional constructs.

In the future, plans are in place to further enhance BiSDL usability by developing a visual language interface and refining the user interface to simplify combining or editing modules. While, in its current implementation, the BiSDL still requires basic programming and modeling skills, a graphical interface would pave the way to the extensive broadening of the user base to people with no programming skills.

Considering the importance of BiSDL adhering to Findability, Accessibility, Interoperability, and Reuse (FAIR) guidelines for data management and stewardship, adoption and enforcement of a standardized naming convention in BiSDL is of paramount goal in the future. In this regard, BiSDL current framework is poised for advancements in two critical areas. Firstly, the versatility of its source-to-source compiler, which currently facilitates the translation of BiSDL into NWN models, could be expanded to support additional formalisms and standards, including those under the COMBINE initiative. Secondly, future works may integrate an ONTOLOGY_ID field into the language within the Metadata representation SBO category to encourage adherence to standards and foster model exchange.

## List of abbreviations

NWN: Nets-Within-Nets
PN: Petri Nets
BiSDL: Biology System Description Language
DBTL: Design–Build–Test–Learn
VHDL: Very High-Speed Integrated Circuit Hardware Description Language
AHL: Acyl-homoserine lactones
3OC6HSL: 3-oxohexanoyl-homoserine lactone
GFP: Green Fluorescent Protein
ACP: acylated acyl carrier protein
SAM: S-adenosylmethionine
LacI: lactose repressor protein
synNotch: synthetic Notch
BFP: blue fluorescent protein
DSL: Domain-Specific Language
GEC: Genetic Engineering of Cells
GUBS: Genomic Unified Behavior Specification
SBOL: Synthetic Biology Open Language
COMBINE: COmputational Modelling in BIology NEtwork
SBGN: Systems Biology Graphical Notation
SBML: Systems Biology Markup Language
ODE: Ordinary Differential Equations
OOP: Object-Oriented Programming
RGB: red, green, and blue
IBL: Infobiotics Language
SED-ML: Simulation Experiment Description Markup Language
gro: The Cell Programming Language
SBO: Systems Biology Ontology
FAIR: Findability, Accessibility, Interoperability, and Reuse
GUI: Graphical User Interface

## Declarations

### Ethics approval and consent to participate

Not applicable

### Consent for publication

Not applicable

### Availability of data and materials

The code and data supporting this study and required to reproduce all published results are publicly available on GitHub at https://github.com/smilies-polito/BiSDL/tree/v1.2 and referenced with Zenodo at https://zenodo.org/doi/10.5281/zenodo.10895847.

### Authors’ contributions

**LG**: Conceptualization, Methodology, Software, Validation, Investigation, Data Curation, Writing - Original Draft, Writing - Review & Editing, Visualization; **RB**: Conceptualization, Methodology, Software, Validation, Investigation, Data Curation, Writing - Original Draft, Writing - Review & Editing, Visualization, Supervision, Funding acquisition; **AS**: Conceptualization, Methodology, Validation, Writing - Review & Editing, Supervision; **SDC**: Conceptualization, Methodology, Validation, Writing - Review & Editing, Supervision, Project administration, Funding acquisition.

## Acknowledgements

Not applicable.

## Authors’ information (optional)

**Leonardo Giannantoni** is a Ph.D. candidate at the Department of Control and Computer Engineering of Politecnico di Torino. He holds an M.Sc. degree in Computer Engineering from Politecnico di Torino in Italy. His research focuses on the application of computational methods to synthetic biology.

**Roberta Bardini** is a post-doc researcher at the Department of Control and Computer Engineering of Politecnico di Torino. She holds a Ph.D. (2019) in Control and Computer Engineering from the Politecnico di Torino in Italy and an M.Sc. in Molecular Biotechnology (2014) from Università di Torino. Her research focuses on computational biology and bioinformatic approaches for analyzing and modeling complex biological systems and on biomanufacturing processes’ computational design and optimization.

**Alessandro Savino** is an Associate Professor in the Department of Control and Computer Engineering at Politecnico di Torino (Italy). He holds a Ph.D. (2009) and an M.S. equivalent (2005) in Computer Engineering and Information Technology from Politecnico di Torino in Italy. Prof. Savino’s research contributions include approximate computing, reliability analysis, safety-critical systems, software-based Self-tests, and image analysis. He has been part of the program and organizing committee of several IEEE and INSTICC conferences and has served as a reviewer of IEEE conferences and journals.

**Stefano Di Carlo** has been a full professor in the Control and Computer Engineering department at Politecnico di Torino (Italy) since 2021. He holds a Ph.D. (2003) and an M.Sc. equivalent (1999) in Computer Engineering and Information Technology from Politecnico di Torino in Italy. Di Carlo’s research contributions include biological network analysis and simulation, machine learning, image processing, evolutionary algorithms, and development of resilient computer architectures.

## Appendix A Basic building blocks

This work proposes a library of BiSDL constructs as an exemplification of its semantic capabilities that show the expressiveness and closeness to the biological semantics of the language. This section illustrates the syntax and semantics of the basic building blocks used to create a BiSDL description, focusing on accessibility and biological significance. Since BiSDL aims to be an easily readable language, it uses English keywords and minimal punctuation. It also avoids cumbersome solutions to delimit blocks. To this end, the use of indentations is suggested, although not mandatory, to set each block apart from the surrounding code visually.

The provided building blocks belong to different domains at Level II, where they provide descriptions of basic biological concepts, including simple local relations in the spatial domain, such as the relative position of two biological elements, and base functions in the behavioral domain, such as the production or degradation of a protein.

### A. Process building blocks

This section provides a detailed overview of the basic constructs offered by BiSDL to model more complex behaviors inside `PROCESS` constructs. All parameters accepted by these building blocks are the base BiSDL types `gene`, `mrna`, `protein`, (protein) `complex`, and generic `molecule`, collectively indicated as *molecules* in the following. All molecule parameters can optionally be associated with a multiplier, i.e., an integer number indicating the ratio between the molecules involved in the process.

**Listing 3.**
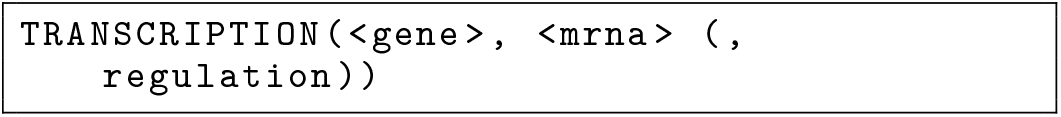
Transcription building block template

The construct in Listing 3 models the copying of DNA segments that can encode proteins in mRNA, the first step in gene expression, with the following parameters:

- `<gene>`: the gene to be transcribed; must end in “_gene”;
- `<mrna>`: the transcribed mRNA; must end in “_mrna”;
- (optional) `regulation` is a list of molecules acting as inhibitors, inducers, or activators (see subsection A-B).

**Listing 4.**
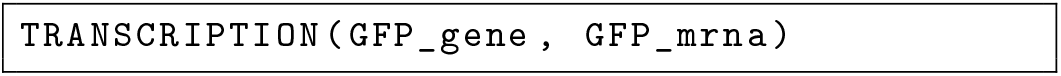
Transcription building block example

Listing 4 provides an example of the transcription of a gene to an mRNA.

**Listing 5.**
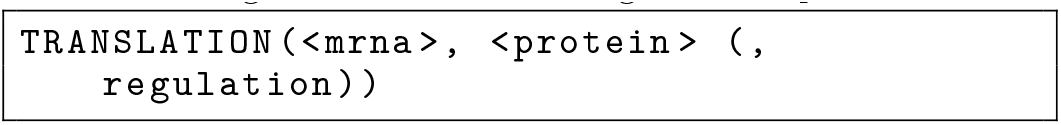
Translation building block template

The construct in Listing 5 models the process of protein synthesis from their mRNA blueprint, with the following parameters:

- `<mrna>`: the mRNA to be translated; must end in “_mrna”;
- `<protein>`: the protein to be produced; must end in “_protein”;
- (optional) `regulation` is a list of molecules acting as inhibitors, inducers, or activators (see subsection A-B).

**Listing 6.**
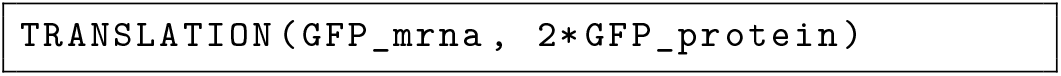
Translation building block example

Listing 6 provides an example of the translation of a mRNA to two proteins.

**Listing 7.**
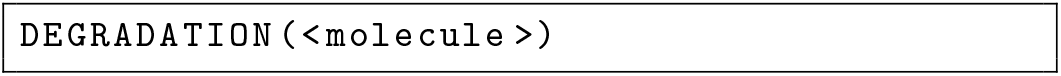
Degradation building block template

The construct in Listing 7 models degradation of a molecule, with the following parameter:

- `<molecule>`: the molecule to be broken down.
- (optional) `regulation` is a list of molecules acting as inhibitors, inducers, or activators (see subsection A-B).

**Listing 8.**
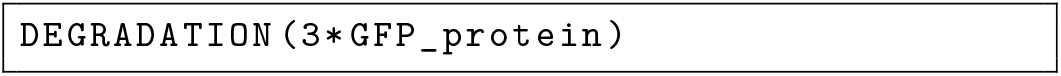
Degradation building block example

Listing 8 provides an example of the degradation of a protein.

**Listing 9.**
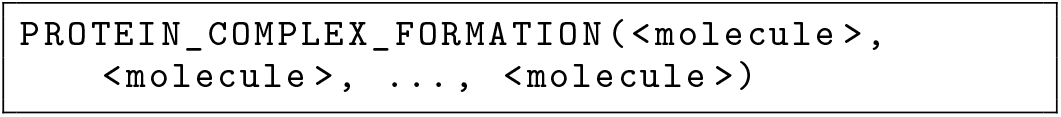
Complex formation building block template

The construct in Listing 9 models the process of two or more proteins associating in a protein complex, with the following parameter:

- `<molecule>, …`: the proteins (at least two) that participate in the formation of the complex, followed by the name of the protein complex itself.

**Listing 10.**
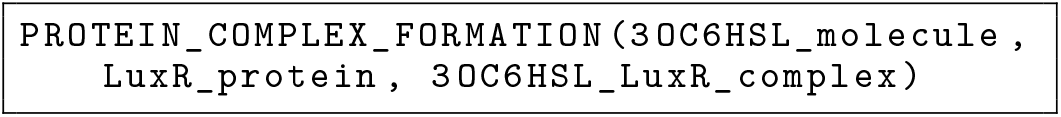
Complex formation building block example

Listing 10 provides an example of the formation of a protein complex.

**Listing 11.**
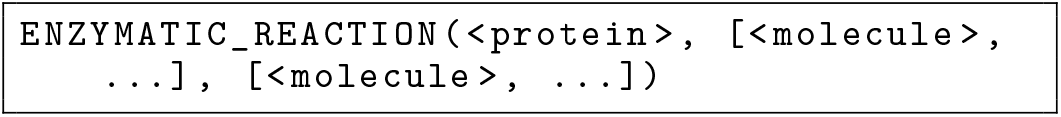
Enzymatic reaction building block template

The construct in Listing 11 models the enzyme catalysis mechanism, through which a catalyst facilitates the reactants in the formation of one or more products, with the parameters:

- `<protein>`: the catalytic enzyme;
- `[<molecule>, …]`: a list of at least one reactant, i.e., the *input* molecules;
- `[<molecule>, …]`: the list of at least one product formed in the process, i.e., the output molecules.

**Listing 12.**
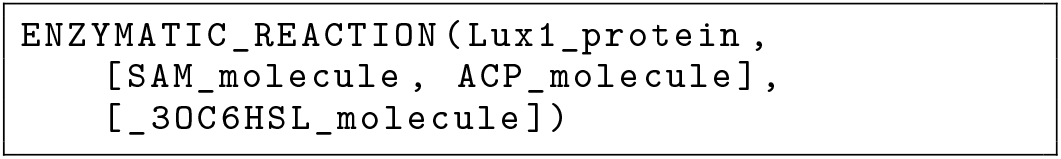
Enzymatic reaction building block example

Listing 12 provides an example of an enzymatic reaction with two reagents and one product.

**Listing 13.**
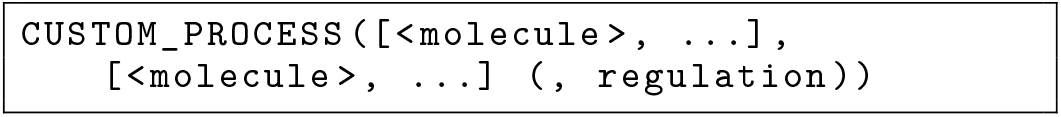
Custom process building block template

The construct in Listing 13 allows the modeling of biological concepts not covered by the previously described ones, with the parameters:

- `[<molecule>, …]`: a list of at least one reactant, i.e., the *input* molecules;
- `[<molecule>, …]`: the list of at least one product formed in the process, i.e., the *output* molecules;
- (optional) `regulation`: a list of molecules acting as inhibitors, inducers, or activators (see subsection A-B).

**Listing 14.**
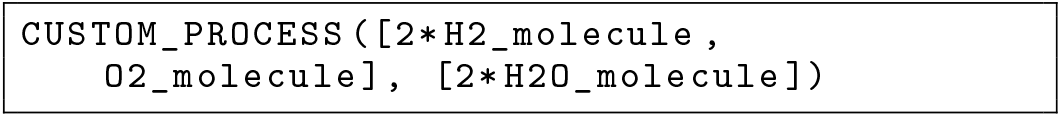
Custom process building block example

Listing 14 provides an example of a custom process.

### B Regulation

It is possible to optionally specify a list of regulatory mediators for the TRANSCRIPTION, TRANSLATION, DEGRADATION, and CUSTOM_PROCESS constructs, to increase or decrease the yield of the process.

**Listing 15.**
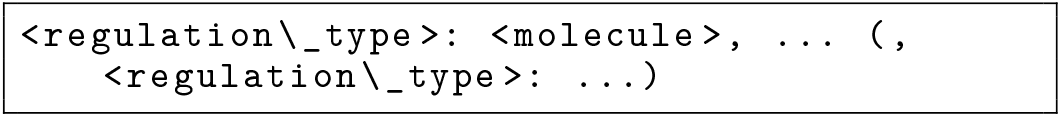
Regulation mechanism template

Listing 15 shows the regulation construct, that accepts the following parameters:

- `<regulation_type>`: `INDUCERS`, `INHIBITORS`, or`ACTIVATORS`;
- `<molecule>, …`: a list of molecules acting as inhibitors, inducers, or activators.

**Listing 16.**
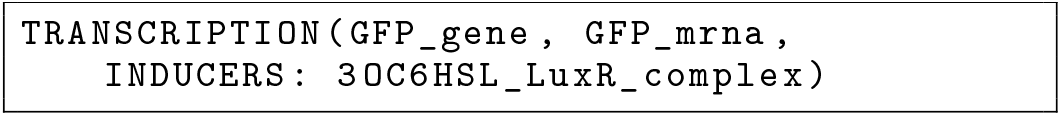
Regulation mechanism example

Listing 16 exemplifies the transcription of a gene to an mRNA, induced by a protein complex.

### C. Signaling mechanisms

The signaling mechanisms available in BiSDL enable the exchange of specific information between `SCOPE` entities:

- from a source entity towards its neighbors;
- from a source entity to a linked destination entity;
- bidirectionally, between two entities.

The first two – implemented by the constructs PARACRINE_SIGNAL and `JUXTACRINE_SIGNAL`, respectively – must be specified inside the `SCOPE` context, as they rely on the properties of the enclosing biological compartment. On the other hand, the specification of the third one (i.e., the `DIFFUSION` construct) must be inside the `MODULE` context after the description of all `SCOPE` statements, as it is a property equally shared between two `SCOPE` statements.

The BiSDL types that can be involved as `signal` parameters are the `protein`, `complex`, and `molecule` types (i.e., `gene`, `mrna`, and `receptor` types are not signals).

**Listing 17.**
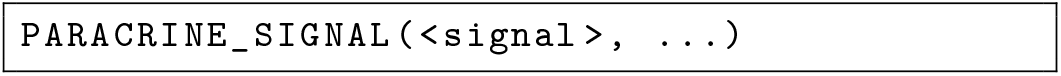
Paracrine signaling template

Listing 17 models the communication mechanism from a cell to its neighboring cells. It requires the specification of a signal (or a list of signals). Spatial coordinates allow for automatically inferring the neighborhood of the container `SCOPE`. The signals reach the destination `SCOPE` unchanged. It accepts the following parameters:

- `<signal>, …`: one or more `protein` or `molecule` entities acting as a signal;

**Listing 18.**
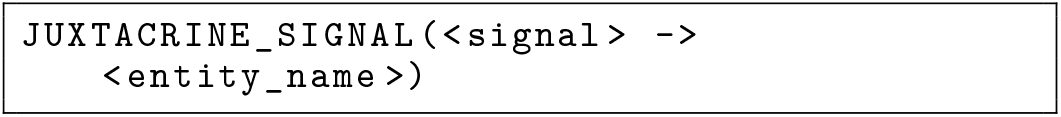
Juxtacrine signaling template

Listing 18 models contact-dependent signaling from a cell to a specific linked cell. It requires the signal and the receiving entity to be specified, with the following parameters:

- `<signal>`: the `protein` or `molecule` entity acting as a signal;
- `<entity_name>`: the destination `SCOPE`.

If the signal is a `molecule` (i.e., its name contains the tag “_molecule”), the corresponding construct models a cellular junction through which the signal is transferred from the source compartment to the destination one (Figure 18). On the other hand, if the signal involved is a `protein` (i.e., its name contains the tag “_protein”), it is interpreted as a membrane ligand and *consumed* in the bond with an active receptor (Figure 19 and Figure 28).

**Fig. 18.**
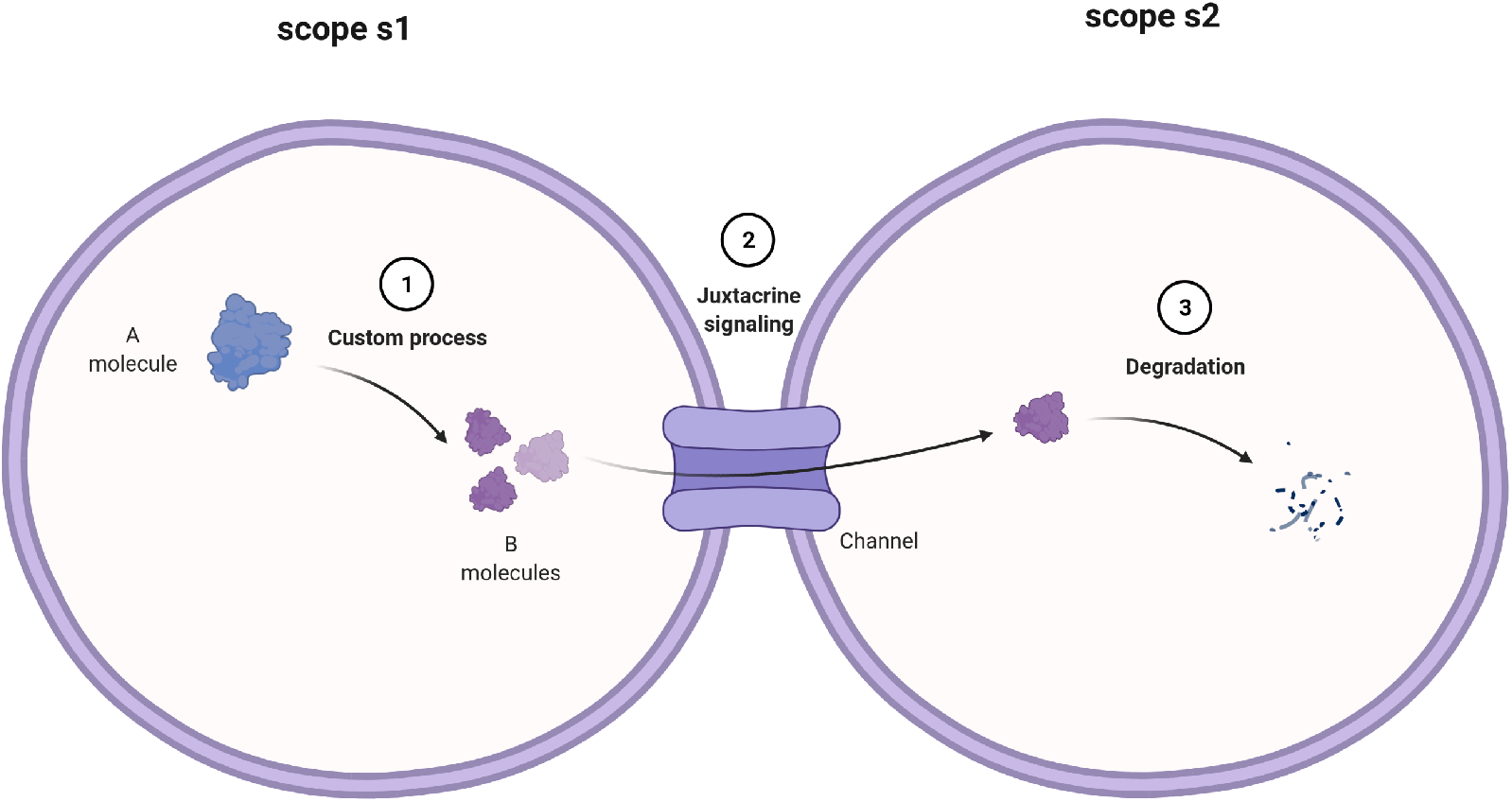
This mock process modeled in Listing 19 involves two biological scopes, `s1` and `s2`. `s1` is the location where three `B_molecule`s are produced (1) starting from one `B_molecule. B_molecule`s are allowed to transit through a communicating junction from scope s1 to scope s2 (2). In `s2, B_molecule`s are constantly degraded (3).

**Fig. 19.**
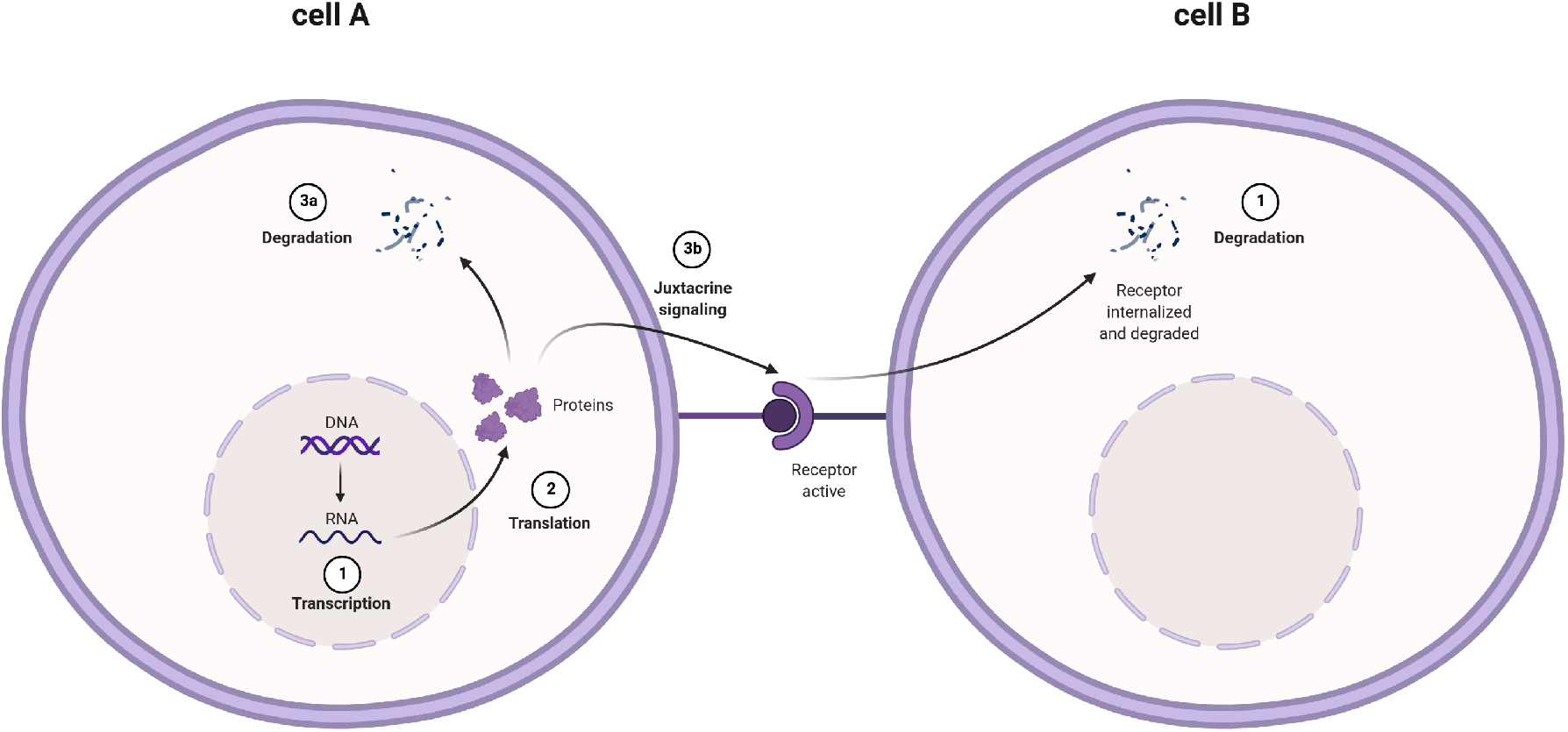
This mock process modeled in Listing 20 involves two biological scopes, `cell_A` and `cell_B. cell_A` is the location where three `A_molecule`s are produced (2) starting from gene-mRNA transcription (1). `A_molecule`s may face degradation (3a), or they could bind (3b) the corresponding receptor on `cell_B` membrane. The active receptor is subject to degradation (`cell_B`, (4)).

As a first example, the BiSDL module in Listing 19 models the mock process depicted in Figure 18 and is compiled to the PN in Figure 27, where a communicating junction links two adjacent cells, allowing the molecule’s transit, where it is modeled with the NWN formalism as follows.

**Listing 19.**
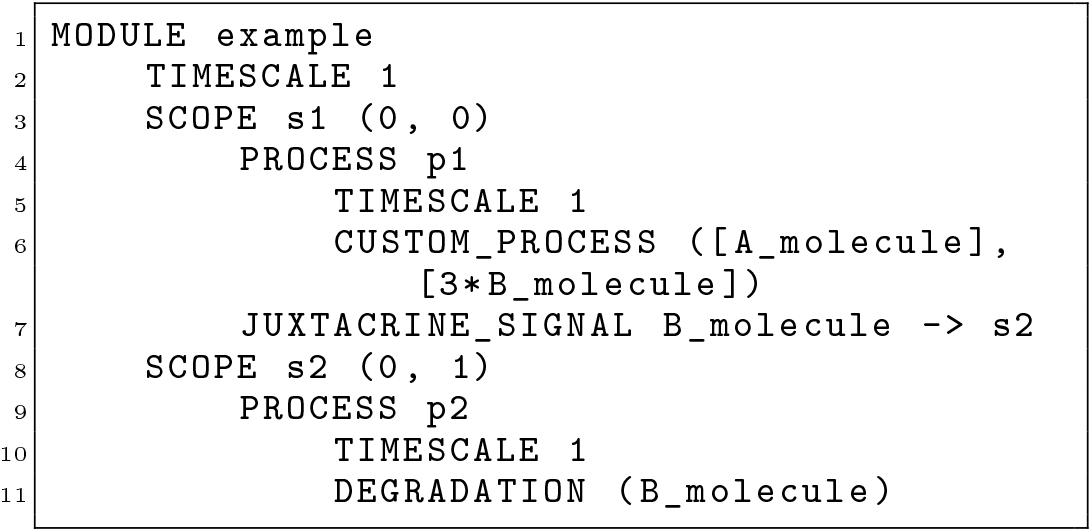
Modeling junction-mediated juxtacrine signaling

As a second example of juxtacrine signaling, Listing 20 models the biological system in Figure 19, where two adjacent cells interact through a membrane ligand (`A_protein`) and a membrane receptor (`A_receptor_active_protein`).

**Listing 20.**
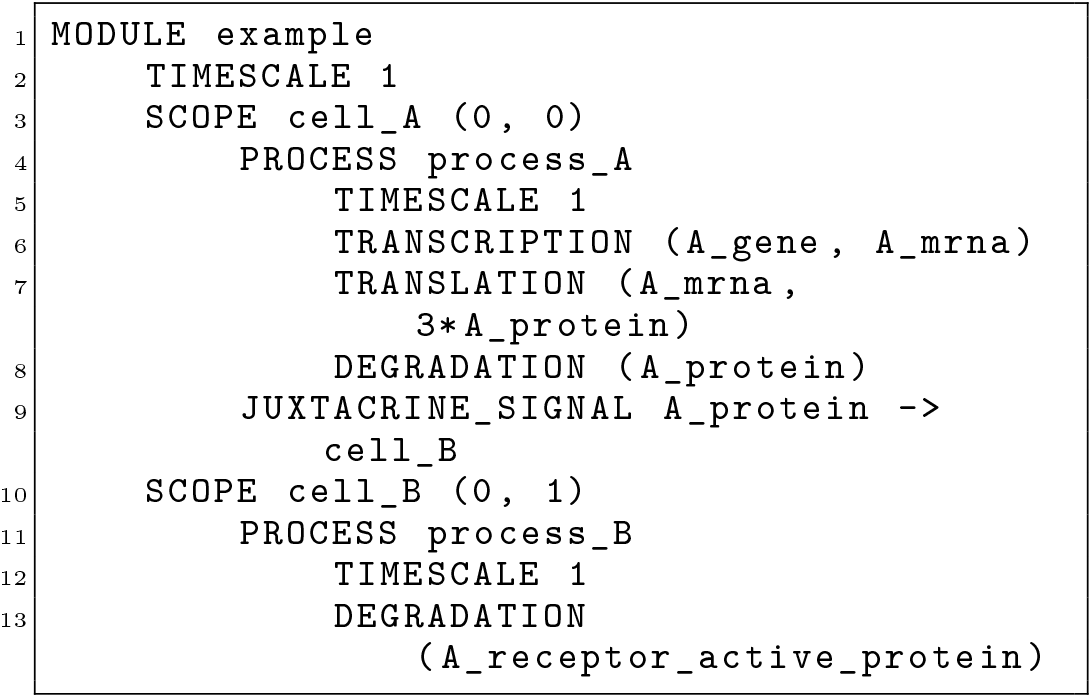
Modeling ligand-receptor juxtacrine signaling

In BiSDL, the declaration of the juxtacrine signaling from `cell_A` to `cell_B` (line 9, Listing 20) using the `A_protein` ligand builds the whole ligand-receptor underlying structure. That is, `SCOPE cell_B` is automatically endowed with the (virtual) permanent presence of a membrane receptor for `A_protein` ligand. When the `juxtacrine_signaling_A_molecule_s1_s2_1` transition (Figure 28.a) fires, one `A_protein` token is consumed in `cell_A` and one `A_receptor_active_protein` token is produced in `cell_B`, with the text substitution signifying the activation of the bond. Therefore, there is no need for additional modeling of the signaling in the description of `SCOPE` `cell_B` behavior (lines 10-13, Listing 20). The `DEGRADATION` process (line 13, Listing 20) models the degradation of ligand-receptor complexes.

**Listing 21.**
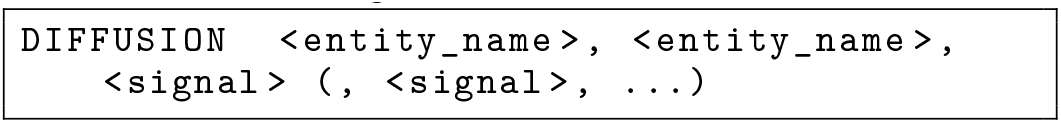
Diffusion template

Listing 21 models a bi-directional permeable membrane between two biological districts, allowing the movement of signals between them, with the following parameters:

- `<entity_name>, <entity_name>`: the two communicating `SCOPE` entities;
- `<signal>, …`: one or more `signal` entities the virtual membrane is permeable to.

**Listing 22.**
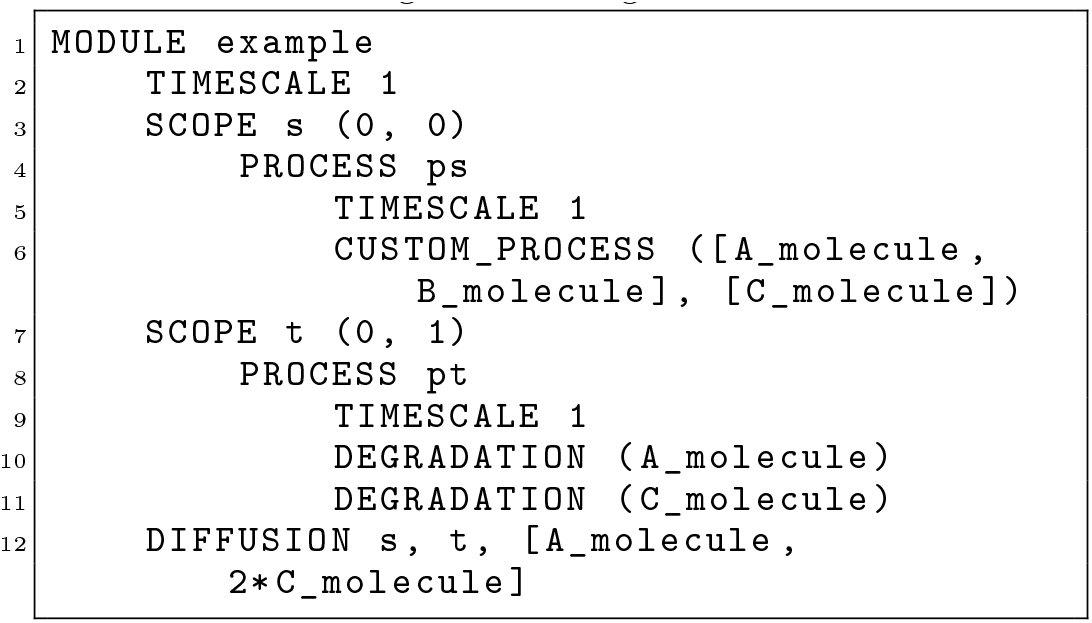
Modeling diffusion

Listing 22 is the BiSDL representation of the biological phenomena in Figure 20. A biological district is divided in two by a membrane. In the leftmost part (a) in Figure 20, a reaction occurs through which `C_molecule` is produced starting from `A_molecule` and `B_molecule`. The membrane is permeable to molecules A and C, with a higher permeability for C. This different permeability is modeled in BiSDL using a multiplier (line 12, Alg. 22).

**Fig. 20.**
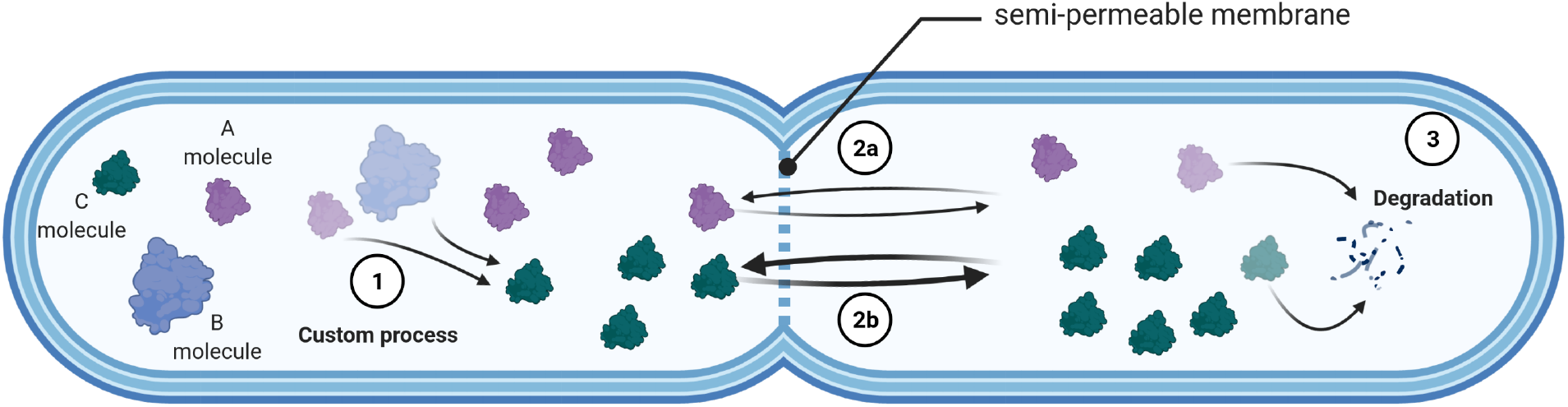
This mock process modeled in Listing 22 involves two biological compartments separated by a semi-permeable membrane allowing the diffusion of `A_molecule` and `C_molecule`. The latter diffuses (2b) twice *as easily* as the former (2a) through the membrane. The compartment left to the membrane is the location where `C_molecule` is produced (1) starting from one `A_molecule` and one `B_molecule`. In the rightmost compartment, the only transformation is the degradation (3) of molecules A and B.

## Appendix B From BiSDL to NWNs

This section details the mapping between BiSDL constructs and the chosen NWN target representation through the `nwn-snakes` library. The `Module` class is a utility layer that eases the simulation task, automatically wrapping the NWN model structure compiled from the BiSDL model description. To facilitate the declaration of variables, the specification of NWN elements in the `Module` class follows the order: PN, `places`, `Transitions`, and finally input and output arcs involving the preceding elements of any level. Each BiSDL `PROCESS` (see Section A-A), once compiled, maps to a net token where places and transitions deal only with black tokens. The black tokens at the lower level correspond, in the top-level *container* place, to colored tokens, i.e., strings bearing the name of the lower-level place holding them. places that are shared among constructs of the same `PROCESS` context are instantiated only once, while their interactions (in terms of arcs and transitions) are created accordingly to the modeled processes.

### A. Compiled process building blocks

As an example, The BiSDL `TRANSLATION` construct in Listing 6 compiles into the PN shown in Figure 21, having one `GFP_mrna` place and one `GFP_protein` place. The two are connected by one `GFP_mrna_translation` transition that, once fired, consumes one token from the `GFP_mrna` place and produces two in the `GFP_protein` place. In this example process, where two proteins (the `(dot, dot)` black tokens) are produced when one mRNA black token (`dot`) is consumed.

**Fig. 21.**
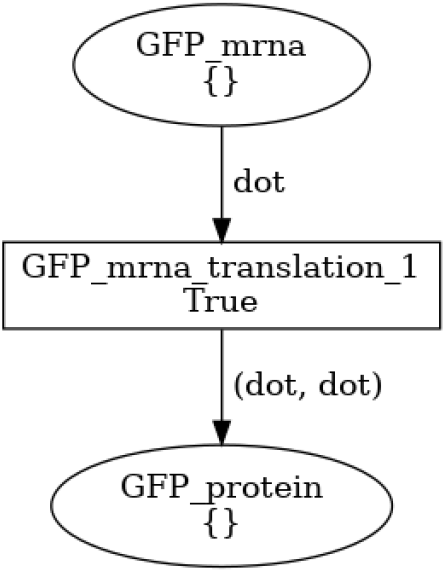
PN for the translation process, generated from the BiSDL code for `TRANSLATION` in Listing 6. Two proteins (the `dot, dot` black tokens) are produced when one mRNA black token is consumed.

As another example, the BiSDL `DEGRADATION` construct in Listing 8 compiles into the PN shown in Figure 22: a `GFP_protein` place is created and connected to three separate transitions that can fire independently, thus depleting up to three tokens.

**Fig. 22.**
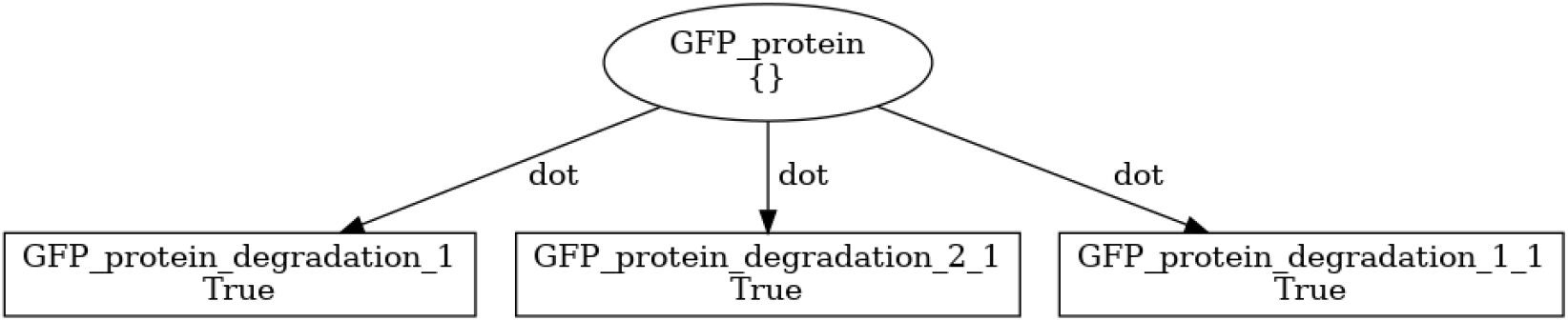
PN for the degradation process corresponding to the construct for `DEGRADATION` in Listing 8. When each one of the transitions fires, one protein is consumed.

For the `ENZYMATIC_REACTION` construct, Listing 12 compiles into the PN shown in Figure 23. In this example, `Lux1_protein` catalyzes the reaction process involving the reactants `SAM_molecule` and `ACP_molecule`, leading to the formation of `3OC6HSL_molecule`. A place is created for the catalytic enzyme and one for each reactant and product. The reactants are input places for the transition representing the enzymatic reaction, which produces one token in the `3OC6HSL_molecule` place upon firing. At the same time, one token is produced back in the `Lux1_protein` place, which, being a catalyst, is not consumed nor changed by the reaction.

**Fig. 23.**
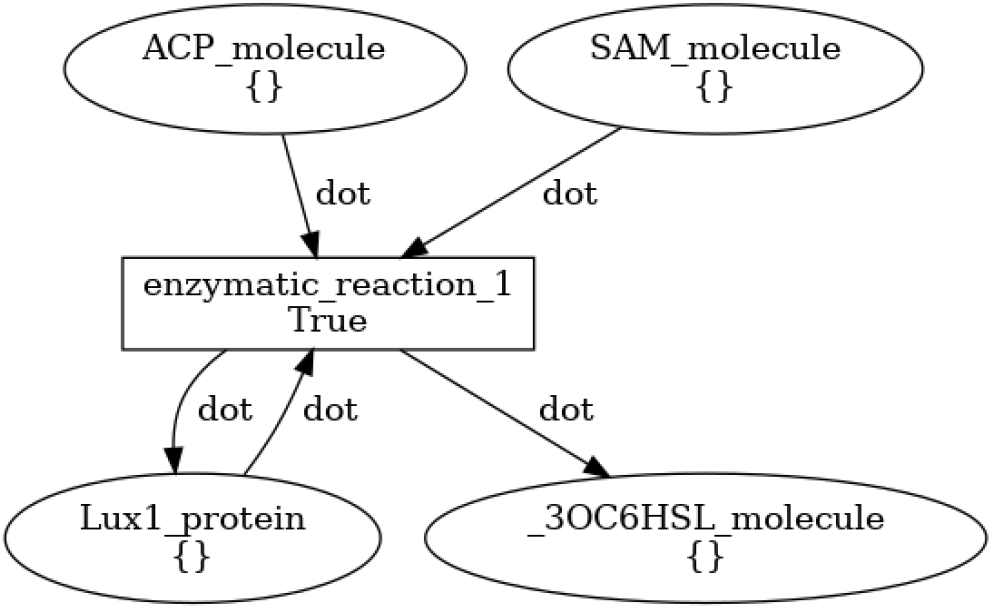
PN for the enzymatic reaction process corresponding to the construct for `ENZYMATIC REACTION` in Listing 12. When the transition fires, one black token is consumed from each of the input places, and one token is produced in the output place (`3OC6HSL_molecule`) and in the place representing the catalysts (`Lux1_protein`).

The BiSDL `CUSTOM_PROCESS` construct in Listing 14 compiles into the PN shown in Figure 24: `H2_molecule` and `O2_molecule` participate in the formation of `H2O_molecule`. One place is created for each molecule. When the transition fires, two black tokens are produced in the `H2O_molecule` place, while two black tokens are depleted from the `H2_molecule` and one from the `O2_molecule`.

**Fig. 24.**
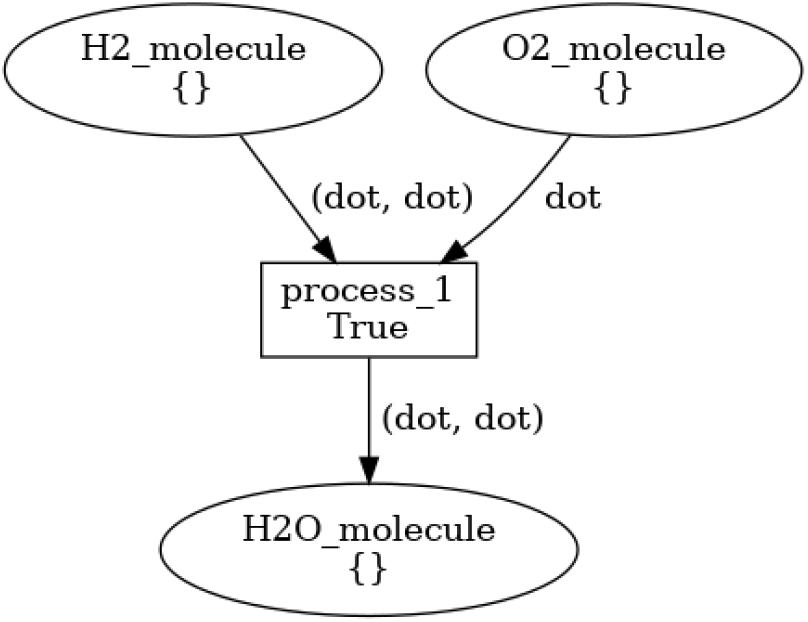
PN for the custom process corresponding to the construct for `CUSTOM PROCESS` in Section Listing 14. When the transition starts, two black tokens are consumed from the `H2_molecule` input places, one from `O2_molecule` input places, and two black tokens are produced in the output place (`H2O_molecule`).

In general, BiSDL descriptions holding these constructs generate a two-level hierarchy of nets. For example, in the context of a complete BiSDL description, the `CUSTOM_PROCESS` construct in Listing 14 generates the described PN model (see Figure 24, and 25.b) at the lower level, and a single place holding it at the top level (Figure 25.a).

**Fig. 25.**
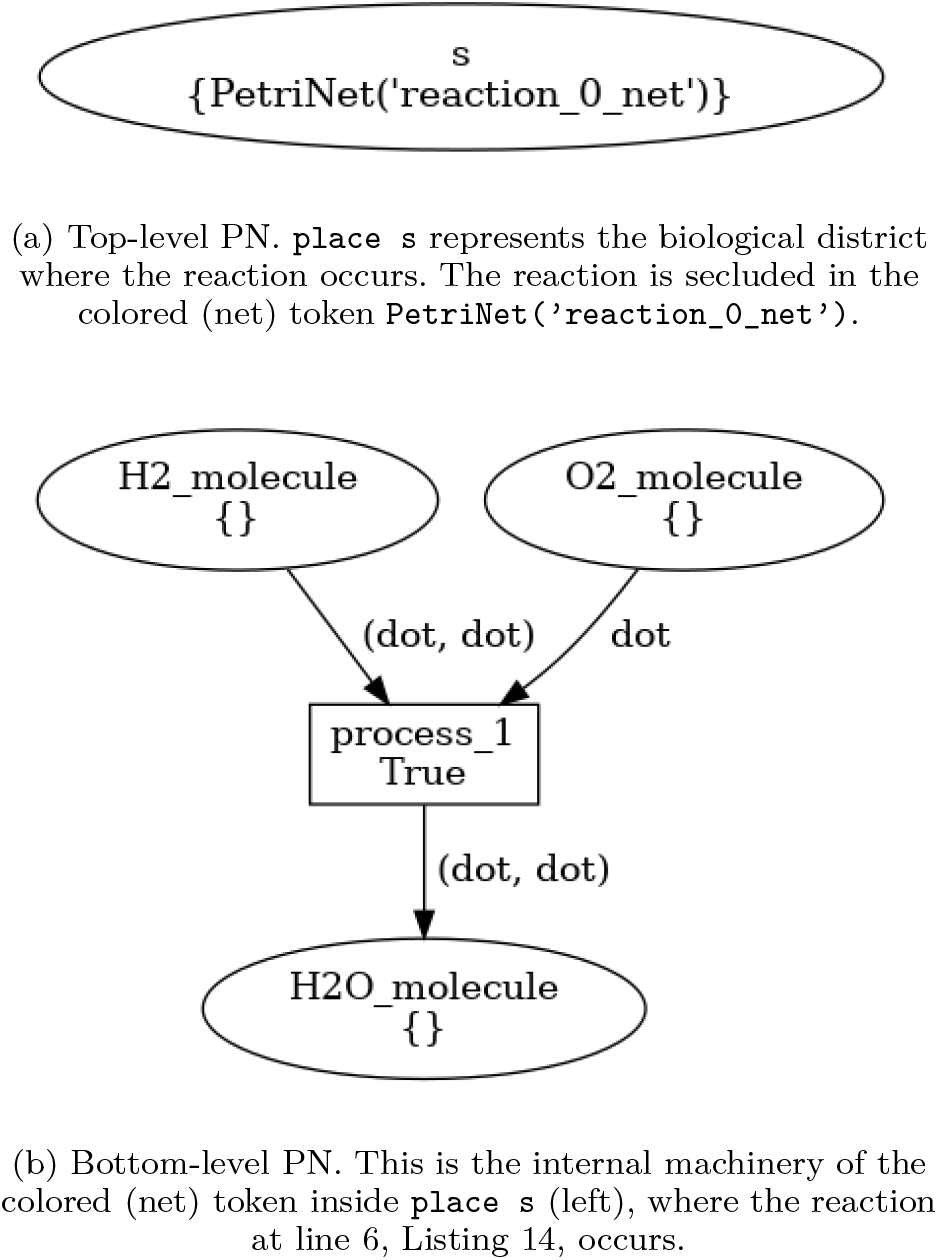
Graphical representation of the PN generated by the compiled BiSDL example in Listing 14.

### B. Compiled regulation constructs

To obtain regulation of basic building block operation, inducer molecules are mapped to additional input places to their main transition. As an example, The BiSDL `TRANSCRIPTION` construct in Listing 16 compiles into the PN shown in Figure 26, including a `GFP_gene` place, a `GFP_mrna` place, and a `3OC6HSL_LuxR_complex` place, connected by one transition. The `GFP_gene_transcription` transition has two input arcs from the `GFP_gene` place and the `3OC6HSL_LuxR_complex` place and two output arcs towards the `GFP_gene` and the `GFP_mrna` place. The gene place is automatically initialized with one black token. The gene token is consumed when the transition fires and is immediately created back. Thus, the net result upon firing is the production of one token in the `GFP_mrna` place and the expenditure of one token in `3OC6HSL_LuxR_complex` place.

**Fig. 26.**
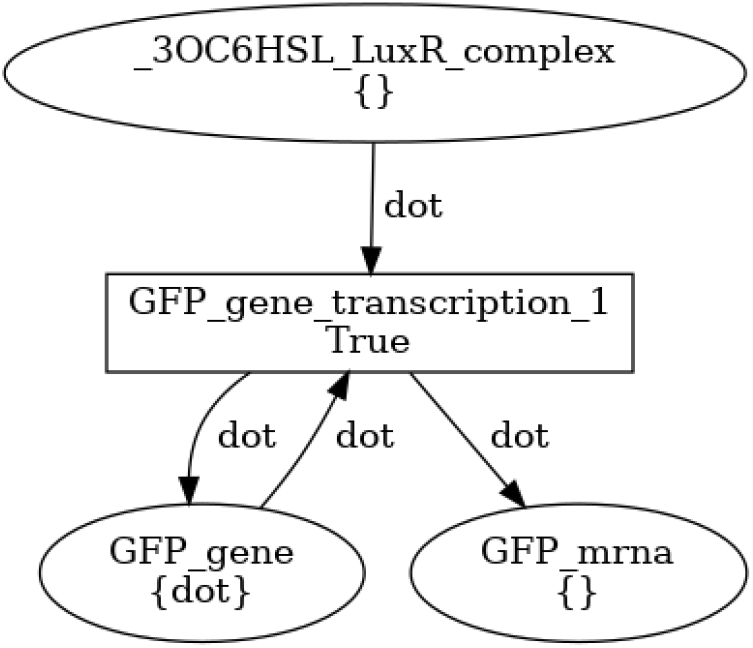
PN for the transcription process, generated from Listing 4, the BiSDL code for `TRANSCRIPTION` in Section A-A.

On the contrary, inhibitory regulation is modeled, at the PN level, with an additional transition competing for the tokens in input to the main transition.

### C. Compiled signaling constructs

The PN structures produced upon compilation of signaling constructs are transitions and arcs that link places in the top-level net (i.e., BiSDL `SCOPE`s).

For example, Listing 19 compiles into the NWN model in Figure 27. In Figure 27.a, `p1_net` is a net token in place `s1`, and `p2_net` is a net token in place `s2`. Their internal machinery is represented in Figure 27.b and Figure 27.c, respectively. `p1_net` and `p2_net` evolution takes place entirely in their respective enclosing places: net tokens do not move across top-level places. Figure 27.b shows the PN `p1_net` corresponding to process p1 (lines 4-6, Listing 19). This is the internal machinery of the colored (net) token inside `place s1` in Figure 27.a. The `process_1` transition gets black tokens from the input place `A_molecule` and produces black tokens in the output place `B_molecule`. Finally, Figure 27.c shows the PN `p2_net`, corresponding to process p2 (lines 9-11, Listing 19), and to the internal machinery of the colored (net) token inside `place s2` in Figure 27.a. The `B_molecule_degradation` transition depletes black tokens from the input place `B_molecule`. Black tokens produced and consumed in `p1_net` places (Figure 27.b) correspond to colored tokens (strings) in the top level `s1` place (Figure 27.a). In general, a low-level black token corresponds to a top-level colored token by the same name of the place it is produced or consumed in. The transition `juxtacrine_signaling_A_molecule_s1_s2_1` is enabled by `A_molecule` colored tokens from `s1`, and outputs the same colored tokens in the output place `s2`. Note that the substitution rule `“protein”, “receptor_active_protein”` has no effect in this example, thus letting `A_molecule` tokens move to the destination scope unchanged. In fact, the biological process we are modeling lets the scopes communicate through a junction (the *Channel* in Figure 18). The scope of this substitution is to model the activation of a receptor when the `JUXTACRINE_SIGNAL` construct is used to model the juxtacrine interaction between a ligand and a membrane receptor, as illustrated in the next example.

The PN structure built after a `DIFFUSION` construct consists, in general, of two transitions per signal. One transition consumes tokens from the first `<entity_name>` and produces the same amount of tokens in the second `<entity_name>`. The other transition works in the opposite direction. When multipliers are specified for a `signal`, the PN structure produced will be composed of as many pairs of transitions as specified by the multiplier.

As an example, Listing 22 compiles into NWN structure in Figure 29. Figure 29.a shows the top-level net structure. places `s` and `t` are mutually connected by one couple of `diffusion_A_molecule_<N>` transitions allowing `“molecule_A”` colored tokens bi-directional passage, and by two couples of `diffusion_C_molecule_<M>` transitions for `“molecule_C”` tokens passage. Such structure allows `C_molecule` to traverse the membrane with twice the probability with respect to `A_molecule`.

**Fig. 27.**
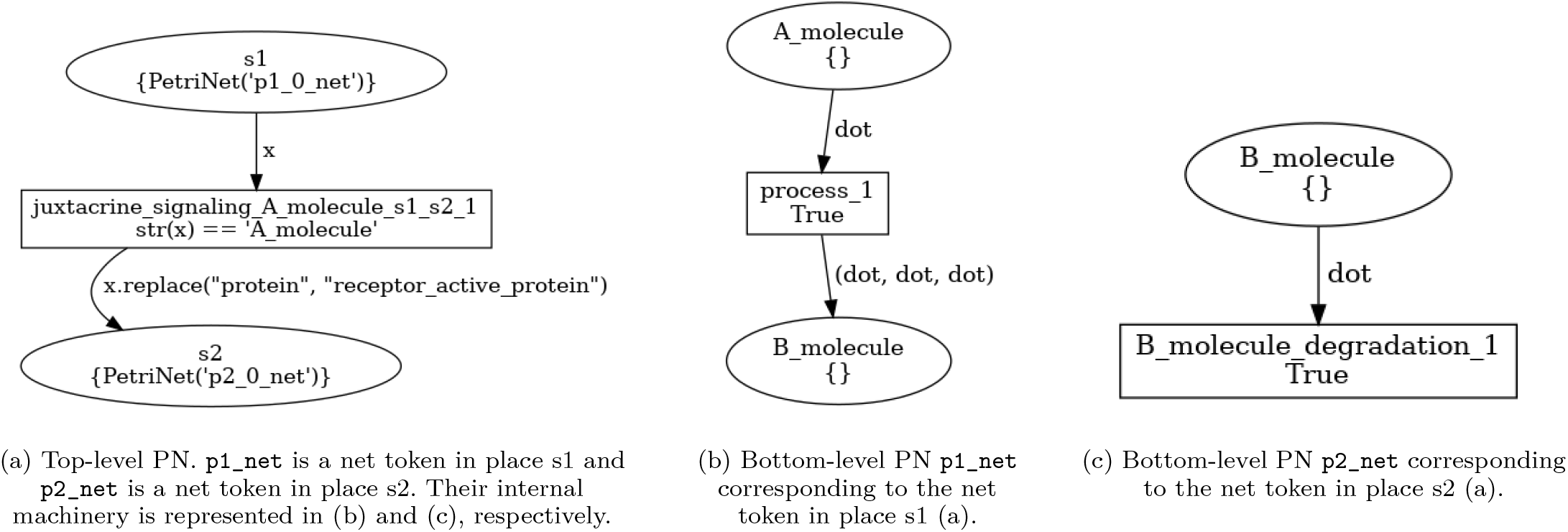
Graphical representation of the PN generated by the compiled BiSDL example in Listing 19.

**Fig. 28.**
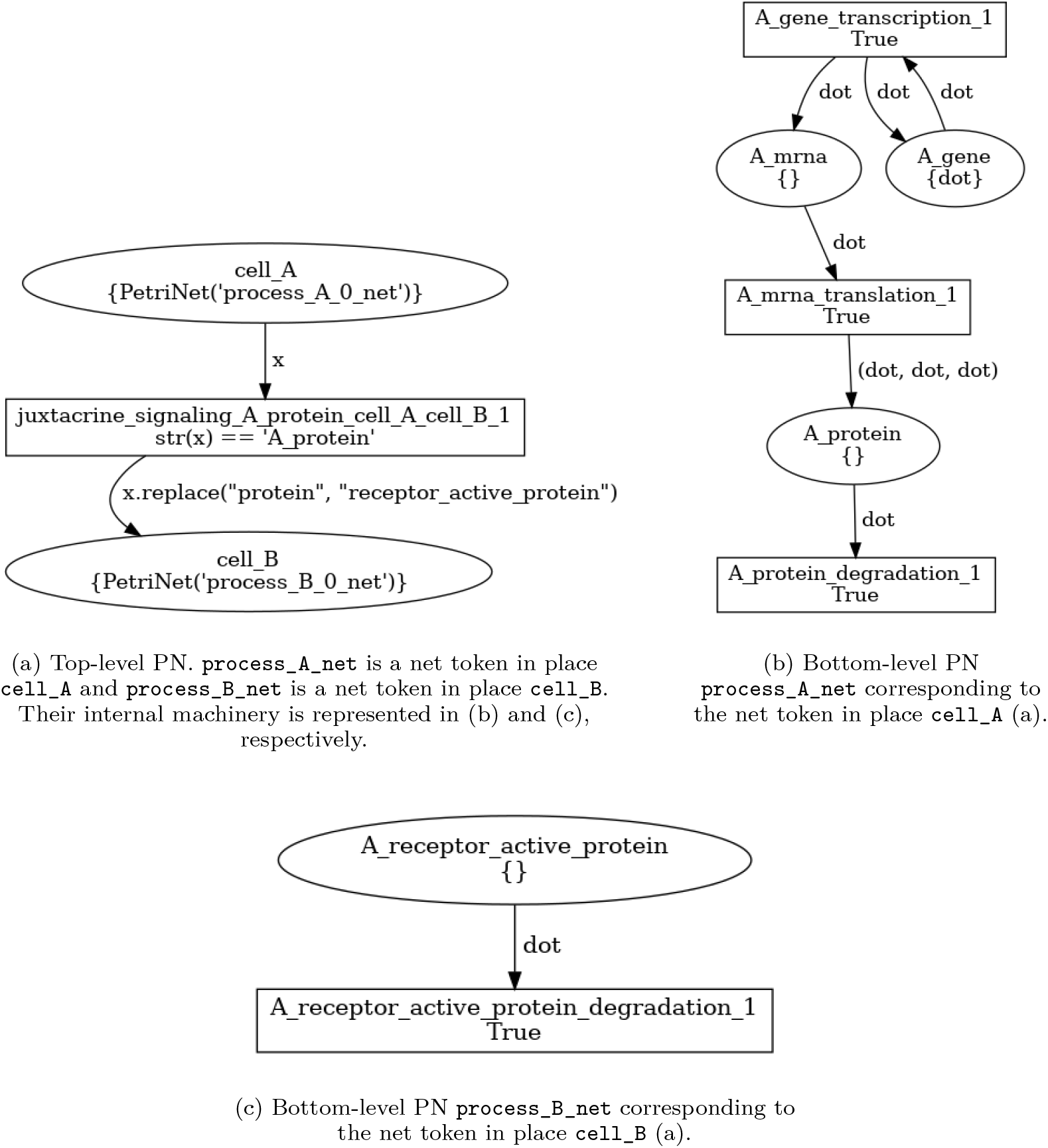
Graphical representation of the PN generated by the compiled BiSDL example in Listing 20.

**Fig. 29.**
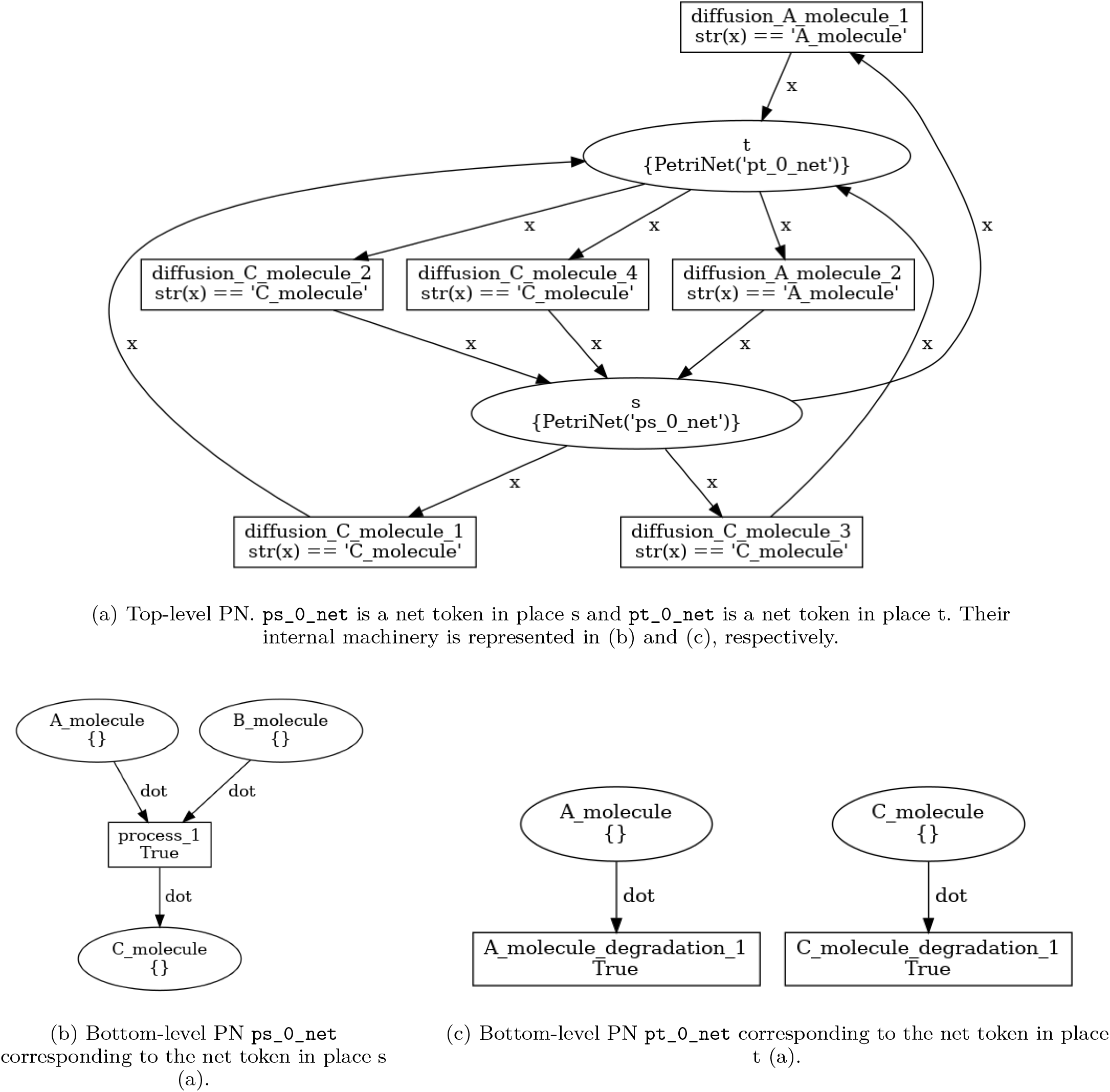
Graphical representation of the PN generated by the compiled BiSDL example in Listing 22.

